# Loss of DHX36/G4R1, a G4 resolvase, drives genome instability and regulates innate immune gene expression in cancer cells

**DOI:** 10.1101/2025.01.03.631217

**Authors:** Anna R. Bartosik, Pei-Chi Hou, James P. Vaughn, Philip J. Smaldino, Aakrosh Ratan, Marty W. Mayo, Wang Yuh-Hwa

## Abstract

G-quadruplexes (G4s) are four-stranded alternative secondary structures formed by guanine-rich nucleic acids and are prevalent across the human genome. G4s are enzymatically resolved using specialized helicases. Previous *in vitro* studies showed that DEAH-box Helicase 36 (DHX36/G4R1/RHAU), has the highest specificity and affinity for G4 structures. Here, by mapping genome-wide DNA double-strand breaks (DSBs), we demonstrate that knockout (KO) of DHX36 helicase increases DSB enrichment at G4 sites and that the presence of the G4 motif is a significant mediator of genome instability at regulatory regions. The loss of DHX36 corresponds with the significant upregulation of NF-κB transcriptional programs, culminating in the production and secretion of proinflammatory cytokines. Loss of DHX36 expression results in an increase in the innate immune signaling stimulator of interferon response cGAMP interactor 1 (*STING1*) expression and activation of genes involved in immune response pathways. Importantly, higher levels of *DHX36* mRNA expression in human B-cell acute lymphoblastic leukemia correlate with improved overall survival relative to lower expression of *DHX36*, highlighting its critical role in preserving genome integrity at a cellular level and in the context of cancer.

## INTRODUCTION

G-quadruplexes (G4s) are noncanonical DNA secondary structures formed by guanine-rich sequences. These structures are characterized by stacked planar arrays of guanine tetrads stabilized through Hoogsteen hydrogen bonding (1–3). The polymorphic nature of G4s allows them to form via intra- or intermolecular interactions, influenced by strand orientation and spacer lengths between guanines. *In vitro* studies using purified single-stranded DNA as a template under G4-promoting conditions have identified over 700,000 G4-induced polymerase-stalling sequences (4, 5), and subsequent genome-wide analyses using a structure-specific antibody (BG4) have validated the presence of a subset of these sequences in human cells (6–10). The abundance and evolutionary conservation of G4 structures across species underscores their functional role at both genomic and transcriptomic levels (5).

G4 structures are enriched in key regulatory regions such as enhancers, promoters, and transcription factor binding sites, where they have been implicated in transcriptional regulation (11–14). G4s have been associated with both repressing and promoting transcription. G4 DNA forms a physical barrier that impedes RNA polymerase progression along the template strand (15, 16). Alternatively, the presence of a G4 structure on the non-template strand can interfere with transcription by suppressing the reannealing of the DNA helix behind the RNA polymerase complex and facilitating the formation of stable DNA:RNA hybrids that will block on-coming transcriptional machinery (17). On the other hand, G4s in promoter regions are associated with enhanced transcriptional activity, serving as recruitment platforms for transcription factors (7, 18, 19). Additionally, the formation of G4s in promoters contributes to the hypomethylation of CpG islands, resulting in elevated gene expression (20). In addition to transcription regulation, G4 structures can also influence DNA replication, by forming the structure on either the leading or lagging strands to sterically block or stall replication forks (21–24). Under these instances, G4 structures can cause DNA double-strand breaks (DSBs), disrupt histone recycling, or increase mutational rate and genome instability (22, 25–27).

Aberrant G4 formation has been linked to nucleotide substitution frequency changes (28) and enrichment of cancer-associated mutations, suggesting a role in promoting somatic mutagenesis (27, 29). Analysis of a large cohort of cancer patients indicates a significant positive correlation between the local density of G4s and single nucleotide variants in cancer patients, suggesting G4-mediated genome instability and the accumulation of somatic mutations (25). Additionally, stabilization of G4s by small ligands results in an accumulation of DNA damage, evidenced by phosphorylation of H2AX and activation of the ATM/ATR pathway (30). However, γH2AX detection and microscopy-based approaches lack the resolution to assess G4-induced DNA damage and its mechanistic implications.

Due to their high thermodynamic stability, G4 structures necessitate enzymatic resolution by specialized helicases to prevent genomic instability (1, 3). Dysregulation or deficiency in G4-DNA helicases can lead to DNA damage, aberrant cell growth, and various diseases (31). Unresolved G4s can promote DSB formation, particularly in cells lacking functional G4-resolving helicases. Impairment of helicases such as BLM, WRN, XPD, FANCJ, or PIF1 results in different pathological syndromes associated with elevated cancer predispositions (32–34). Among multiple G4 helicases, DHX36, also known as RHAU or G4R1, a member of the DExD/H family, resolves both DNA and RNA G4 with high affinity (35–38). *In vitro* studies showed DHX36 has a low nanomolar affinity for DNA G4 structures and has the highest specificity for G4s among DEAH-box helicases (39, 40). Moreover, DHX36 gene silencing increases the level of G4 structures within cells (41, 42). DHX36 selectively binds and unwinds G4s (35, 36, 39, 40, 43–45) and is essential for heart development, hematopoiesis, and embryogenesis in mice (46–48). Thus, the requirement for DHX36 strongly supports the importance of DHX36 in mouse development and highlights the non-redundant function of the DHX36 helicase activity.

In this study, we investigate the role of DHX36 in maintaining genome stability using human T-lymphoblastoid Jurkat cells as a model system. Our findings indicate that the DHX36 loss results in elevated levels of DNA DSBs enriched at G4-containing regions. Specifically, we show that regulatory regions, such as transcription start sites, H3K27ac, H3K4me3, and CCCTC-binding factor (CTCF)-binding sites, when enriched with G4, exhibit significantly higher levels of DNA DSBs. Furthermore, transcriptomic analysis of DHX36-deficient cells uncovered global changes in gene expression, with enrichment of NFκB-regulated immune gene signatures. Genes significantly upregulated upon DHX36 loss were predominantly involved in immune response pathways and known targets of the NF-κB transcription factor. DHX36 depletion also activated the NF-κB signaling pathway, accompanied by enhanced production of proinflammatory cytokines. Collectively, our findings highlight the critical role of DHX36 helicase in preserving genome integrity and regulating immune signaling pathways. Notably, high expression levels of DHX36 correlate with improved survival and prognosis in pediatric B-acute lymphoblastic leukemia (B-ALL) patients, further emphasizing its clinical relevance.

## MATERIALS AND METHODS

### Cell culture

Two Jurkat (GenScript) DHX36 knockout cell lines (KO1 and KO2) were generated by the CRISPR/Cas9 method with the same gRNA (AAGTACGATATGACTAACAC) that targeted the exon 8 region of the *DHX36* gene and resulted in different frameshift mutations in the same exon (49, 50). Cells were cultured at 37°C, 5% CO_2_ in RPMI 1640 medium (Gibco, 11875093) supplemented with 10% fetal bovine serum (FBS, Gibco, 16000044). The parental cell line (wild type, WT) was verified using the short tandem repeat profiling and showed a 98% match to the ATCC database profile (ATCC). Cells were tested negative for mycoplasma using the LookOut Mycoplasma qPCR Detection Kit (Sigma).

### Protein fractionation and western blot

Protein fractionation, as previously described (51), was performed with Jurkat WT, KO1, KO2, and WT treated with human tumor necrosis factor (TNFα, Gold Biotechnology, cat: 1130-01-10) for 17min. Western blotting of whole cell extract, cytoplasm and nuclear fractions was performed as previously described (52) with primary antibodies (DHX36 [Santa Cruz, 1:1000, cat: sc-377485], GAPDH [Cell Signaling, 1:10,000, cat: 5174], CTCF [BD Fisher, 1:1,000, cat:612149], p65 [Cell Signaling, 1:1000, cat:8242], and STING [Cell Signaling, 1:1000, cat:13647]. For γH2AX (Cell Signaling, cat: 9718) detection, cells were lysed in lysis buffer (10mM Tris-HCl, pH=8.0, 1% Triton X-100, 0.1% sodium deoxycholate, 0.1% SDS, 2mM MgCl_2_, 1X protease inhibitors [cOmplete mini EDTA-free, Roche], and 1X phosphatase inhibitors [PhosStop, Roche]) for 15 min at 4°C followed by 15 min incubation with Benzonease nuclease (50U/mL, Sigma cat: 70746-3), and western blotting was performed.

### MTS proliferation assay

Jurkat WT, KO1, and KO2 cells were seeded in a 96-well plate and analyzed using CellTiter96^®^ AQueous One Solution Cell Proliferation Assay (Promega, cat. G3580) following the manufacturer’s protocol.

### Reverse-transcription and real-time RT-PCR

Total RNA was isolated from cell pellets using Trizol reagent (Life Technologies, cat. 15596018) according to the manufacturer’s protocol. Reverse-transcription reaction was performed using kit (SuperScript™ IV First-Strand Synthesis System, cat: 18091050) following the manufacturer’s protocol. BioRad CFX384 Touch Real-Time PCR Detection System was used for real-time polymerase chain reaction (PCR). To avoid genomic DNA contamination, all primers were designed to span cross-exon junctions, and the expression level for each gene was normalized to the *GAPDH*. Primer sequences are included in **Supplementary Table S1.**

### RNA sequencing

Total RNA from Jurkat WT and DHX36 KO cells was extracted and enriched for poly-A fragments, followed by fragmentation, reverse transcription, and second-strand complementary DNA (cDNA) synthesis. Double-stranded cDNA was purified with AMPure XP beads, and ends were repaired and then ligated with Illumina sequencing adapters and PCR amplified (Novogene, Inc.). Libraries were then subjected to 150-bp paired-end reads sequencing on the NovaSeq6000 Illumina platform. Jurkat WT RNA-seq was previously published (52).

### Luminex^®^ Multiplex Assay

For the cytokines array assay, Jurkat cells WT, KO1, and KO2 were seeded, and 24 h later, the supernatant, along with the cell-free medium, was collected. The supernatant was subsequently used for Luminex® Multiplex Assay ELISA Human 47 Plex Panel and Human Cytokine/Chemokine Panel II assays. Targeted cytokine levels upon loss of DHX36 were normalized to a cell-free medium and calculated as fold change compared to WT.

### Genome-wide break mapping and sequencing

For DNA double-strand break detection, genomic DNA was purified by lysing cells in lysis buffer (0.5M Tris-HCl pH 8.0, 0.1M EDTA, 0.1M NaCl, 1% SDS, 1mg/mL Proteinase K) for 3h at 55°C followed by organic extraction and ethanol precipitation. Gentle pipetting with wide-open pipette tips was used to avoid DNA shearing and introduction of artificial DNA breaks during purification. Detection of DSBs was performed as previously described (53–56). Briefly, genomic DNA was subjected to blunting/A-tailing reactions followed by Illumina P5 adaptor ligation to capture broken DNA ends. P5-ligated DNA fragments were then purified and fragmented by sonication and subsequently ligated to the Illumina P7 adaptor. Libraries were PCR-amplified and subjected to whole-genome 150-bp paired-end sequencing with the Illumina HiSeq X Ten platform.

### Processing of DSB reads

Sequencing reads were aligned to the human genome (GRCh38/hg38) with the bowtie2 (v. 2.3.4.1) aligner running in high sensitivity mode (--very-sensitive). The fragment length was restricted from 100 nt to 2000 nt (-X 2000 -I 100 options). Unmapped, non-primary, supplementary, and low-quality reads were filtered out with SAMtools (v. 1.7) (-F 2820). Further, PCR duplicates were marked with Picard-tools (v. 1.95) MarkDuplicates, and finally, the first mate of non-duplicate pairs (-f 67 -F 1024) was selected with SAMtools for downstream analysis. For each detected break, the most 5’ nucleotide of the first mate defined the DNA break position. Sequencing and alignment statistics for the DSB mapping/sequencing libraries prepared from Jurkat DHX36 KO cells are listed in **Supplementary Table S2**. Jurkat WT DNA DSB mapping sequencing data was previously published (52). Biological replicates showed high reproducibility of genomic coverage (Pearson’s correlation r = 0.922-0.989, p ≅ 0, for KO1; and r = 0.974-0.995, p ≅ 0, for KO2, **Supplementary Figure S1,** r=0.926-0.989 for WT(52)). and were subsequently combined for downstream data analysis. This strong correlation confirms that the break mapping procedure does not introduce significant amounts of random DNA breaks that could convert single-stranded nicks into DSBs.

### Peak calling

The macs2 software tool (v. 2.2.9.1) was used to call peaks (57). For chromatin immunoprecipitation assays with sequencing (ChIP-seq) data, macs2 was run with default settings, and each dataset was controlled for the matching input control (-c). For peak calling in DSB data, the blacklist region was removed before peak calling. No input data was used for Jurkat WT and DHX36 KO samples, and a no-shift model was employed because the break data is defined by the 5’ end of read1 alone. Called peaks from WT, KO1, and KO2 were merged using BEDTools (v. 2.27.1), and a total of 33 304 unique break cluster sites, which represent 0.43% of the human genome, were used for the average plots, heatmaps, and box plots.

### Identifying consensus G4-forming regions

To generate G4 consensus sites, publicly available BG4 ChIP-seq data from NHEK, K562, HaCaT, and NSC cells and CUT&Tag data from U2OS and HeLa cells were downloaded through Sequence Read Archive (**Supplementary Table S3**) and aligned to the human GRCh38/hg38 genome using bowtie2 (v. 2.3.4.1). Biological replicates were merged. Low-quality, secondary, and supplementary alignments and unmapped reads were filtered out. Peaks were called by macs2 (v. 2.2.9.1) for each cell line controlled by input or IgG control (-c). Using BEDTools (v. 2.27.1), a merged union list of BG4 binding sites was created (n = 77 125) and used for intersection and reporting the number of peaks that were present at the same genomic location in at least five out of six cell lines; those sites were extracted and defined as G4 consensus sites (n = 4 880). To generate G4 random shuffle control, BEDTools (v. 2.27.1) *shuffle* was used to randomly choose genomic locations and keep features of G4 consensus sites on the same chromosome (-chrom).

### Detection of G4 motifs

To detect canonical G4 motifs (G_≥3_ N_1-7_G_≥3_ N_1-7_G_≥3_ N_1-7_ G_≥3_ N_1-7_) located within break cluster sites, regions marked by H3K27ac, H3K4me3, CTCF-binding sites, and transcription start sites (TSSs) spanning +/250bp from the start of the transcription, the non-B DNA database (58) was employed using default settings. For Figure 3, the transcription regulatory regions were binned into five groups based on the gene expression of the WT (for TSS sites) or on the abundance via macs2 score (for H3K27ac, H3K4me3, and CTCF-binding sites). Then, G4 motifs were identified by the non-B DNA database in these binned groups.

### Processing RNA-seq and differential gene expression analysis

Sequencing reads were quantified against a reference GRCh38/hg38 transcriptome downloaded from Ensembl using kallisto (v. 0.48.0) (59). The transcript-level counts were collapsed to gene-level counts using a tximport. Differential gene expression analysis between WT vs. KO1 and WT vs. KO2 was completed using DESeq2 (v. 1.42.1) and R (v. 4.1.2) using a design that accounted for batches. The threshold for significance was set at false discovery rate (FDR) < 0.05 and absolute log-fold change (log_2_FC) ≥ 1 in all conditions. For enriched gene ontology (GO) biological processes, the analysis was done using ShinyGO webtool (v. 0.81) (47) on significantly upregulated and downregulated genes with all expressed genes as background (DE, n = 15 081). We used an FDR cut of 0.05 and selected the ‘Remove redundancy’ option, which outputs the most significant pathway to represent similar pathways sharing 95% of genes. The top pathways were then selected by FDR and sorted based on Fold Enrichment. EnhancedVolcano (v. 1.12.0) R package was used to generate volcano plots for each comparison (60).

### Co-expression analysis

CEMiTool (v. 1.18.1) R package was used (61) for co-expression analysis. A directed signed network was generated using Spearman correlation as the distance metric. Similar modules were merged, and enrichment analysis was performed against ‘GO: Biological Processes 2023’ and ‘ChEA 2022’ libraries downloaded from Enrichr to identify the enrichment of biological themes and transcription factors (TFs) with target genes in the identified modules.

### Generation of NF-κB target gene list

The NF-κB target gene list was composed as follows: Genes and gene products were obtained from Gene Ontology (https://geneontology.org/) using ‘NF-kappaB’ as the search term. To identify evolutionary conserved NF-κB targets, gene lists for *Homo sapiens* and *Mus musculus* were combined, duplicates removed, and mouse-specific genes removed. In parallel, the molecular signatures database (MSigDB, https://www.gsea-msigdb.org/gsea/index.jsp) was used to obtain *Homo sapiens* gene lists for TF targets encompassing NF-κB. Gene lists obtained from GO and MSigDB were combined, and duplicates were removed to generate master lists for classical and non-canonical NF-κB target genes.

### Analysis of B-ALL data

The gene expression counts and clinical data for TARGET-ALL-P2 were downloaded from Genomic Data Commons (GDC) using TCGAbiolinks. Only the bone marrow samples were considered in this study. R packages, survival (v. 3.5-8), and survminer (v. 0.4.9) were used to analyze and generate the survival Kaplan-Meier (KM) plots. This difference in survival remains significant in a Cox-proportional hazards model after accounting for age at diagnosis and gender, which are known to affect survival in B-ALL. DESeq2 was used to identify differences in gene expression between the top 33% of the patients and the bottom 33% of the patients based on the expression of *DHX36*, and EnhancedVolcano was used to generate volcano plots. R package fgsea (v. 1.28.0) was used for enrichment analysis, and xCell (v. 1.1.0) was used for deconvolution analysis.

### Heatmaps and average plots

Heatmaps and accompanying average plots were generated using ngs.plot.r (v. 2.61) (62). For heatmaps for the union break cluster sites (n = 33 304) and G4 consensus sites (n = 4 880), the order of the heatmap was set to be based on the wild type of sample strength of coverage in each region (-GO total), and the scale was set to be consistent across all sample sets (-SC global).

### Correlation plots

The human genome build GRCh38/hg38 was binned into 100 kb windows using BEDTools (v. 2.27.1) makewindows tool. The coverage in each 100 kb bin was calculated for the bam file of each replicate using BEDTools coverage (n = 30 895). The read coverage in each bin was normalized to the total read number (reads per million, PRM). The absolute difference in coverage in each bin between replicates was calculated, and the top 0.05% most different were removed. Bins where both replicates showed zero coverage were removed. Data was read into python3, read normalized read coverage was plotted between two replicates, and Pearson correlation was calculated.

### Genomic region annotation

To assign genomic annotations, BEDTools intersect was used to sequentially assign genomic features, with each region only being assigned to one genomic feature. The sequential feature assignment filters out regions as they are assigned to a feature. The order for assigning genomic features was TSS, promoter, transcription termination site (TTS), and gene body (GB), and those not assigned to any of these features are coded as intergenic. The definitions used for each genomic feature are as follows: promoter region ranging from TSS −1000 nt to −250 nt, TSS region ranging from TSS −250 nt to +250 nt, gene body region ranging from TSS +250 nt to TTS −250 nt, and TTS regions ranging from TTS −250 nt to +250 nt.

### Statistical analysis

Statistical tests were performed using phyton3 (v. 3.6.5) with scipy stats (v. 0.19.1) and R (4.1.2). Tests are specified in figure legends, and asterisks denote statistical significance: * *p* < 0.05, ** *p* < 0.01, *** *p* < 0.001, and **** *p* ∼ 0, unless stated otherwise.

## RESULTS

### Loss of DHX36 helicase results in an increased level of DNA DSBs

DHX36 is a multifunctional helicase involved in various biological processes (31, 63). While DHX36 is responsible for most tetramolecular G4-DNA resolvase activity (35, 36) and is known to affect telomere length (40, 44, 64), its role in regulating global genome integrity has only begun to be elucidated (42). To investigate genome instability with a focus on DHX36 helicase in DNA G4-dependent regulation, we employed a loss-of-function approach by knocking out DHX36 helicase using CRISPR/Cas9 editing with a single gRNA targeting exon 8 in the human T-lymphoblastoid Jurkat cell line. Two Jurkat DHX36 knockout clones were generated (KO1 and KO2) and validated at both protein and RNA levels relative to the Jurkat wild-type parental cell line by western blot and RT-qPCR analysis. Western blot showed a significant reduction of DHX36 protein level by 85% and 88% in KO1 and KO2, respectively (** *p* < 0.01, Student’s t-test, **Supplementary Figure S2A).** RT-qPCR showed corresponding decreases in mRNA levels to 74% and 54% in KO1 and KO2, respectively (* *p* < 0.05, ** *p* < 0.01, Student’s t-test, **Supplementary Figure S2B)**. Despite the loss of DHX36, cell proliferation was not significantly affected (**Supplementary Figure S2C)**. Interestingly, DHX36 KO cells showed an increase in the level of γH2AX detected by western blot (** *p* < 0.01, Student’s t-test, **Supplementary Figure S3)**, indicative of elevated DNA damage. To map and quantify genome-wide DNA DSBs, we used high-resolution DSB mapping/sequencing in both wild-type (WT) and DHX36 KO cells (53–56). Genome-wide DSBs at single nucleotide resolution were mapped, and regions with significant accumulation of DNA breaks (break cluster sites) were identified in each cell line. We identified a higher number of break cluster sites in DHX36 KO1 and KO2 cells compared to WT cells (371 sites in WT, 19 523 sites in KO1, and 23 831 sites in KO2) and observed that most WT break cluster sites overlapped with DHX36 KO break cluster sites (**Figure 1A**). Analysis of 33 304 merged break cluster sites revealed that WT Jurkat cells already displayed elevated DSB levels at these loci, which were further exacerbated in DHX36 KO cells (**\*\*\*\*** *p* ∼ 0, Wilcoxon signed-rank test, **Figure 1B** and **1C**). These observations provide evidence that select regions in the genome are inherently prone to DNA breakage in WT cells, and DHX36 loss exacerbates these vulnerabilities (**Figure 1D**), highlighting the importance of this helicase in maintaining genome integrity. Further examination revealed that the break clusters were preferentially located at transcription start sites (**Figure 1E**). This underscores the essential role of DHX36 helicase in preserving genome integrity, particularly within transcriptionally active regions.

**Figure 1.**
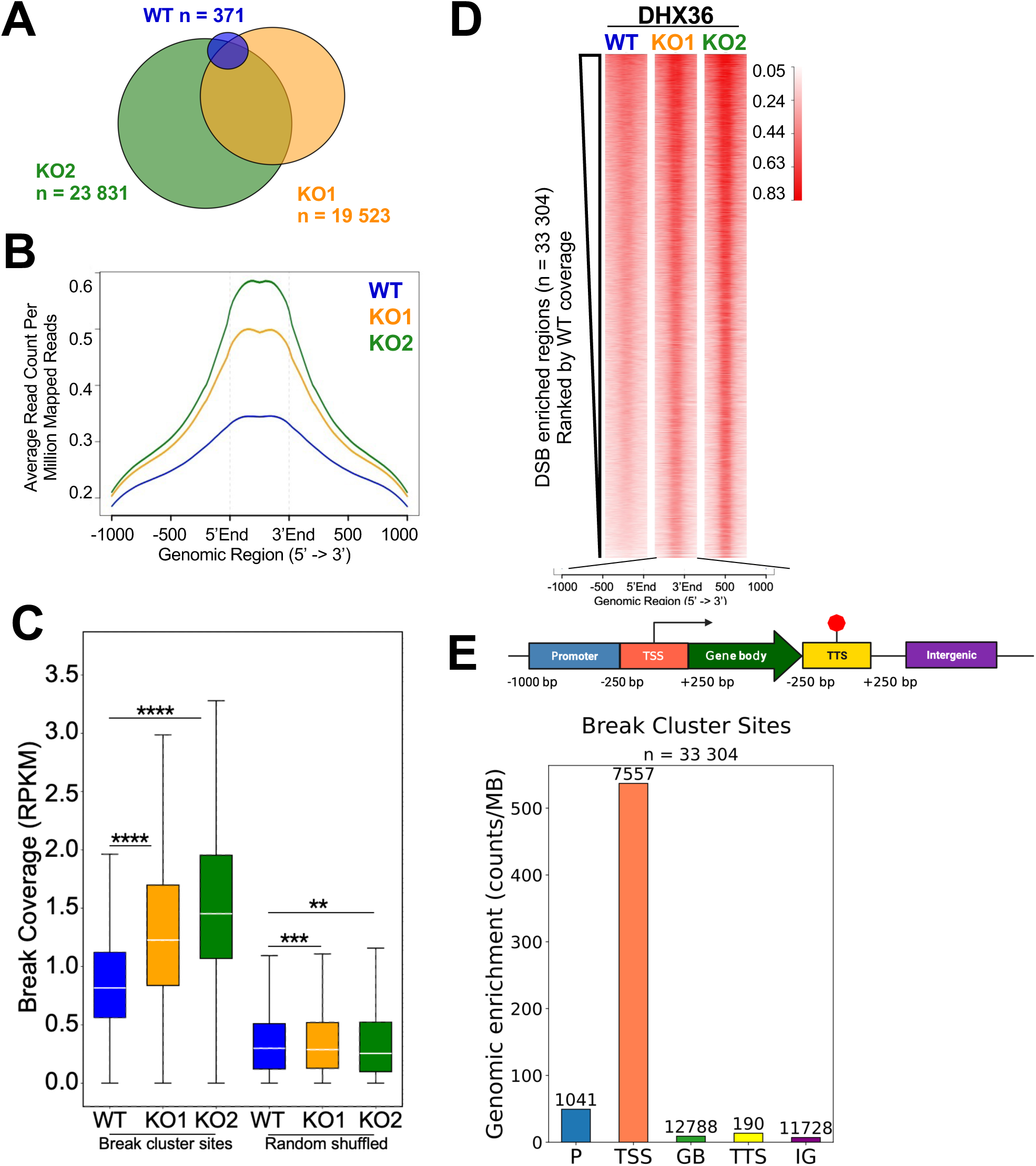
Depletion of DHX36 helicase results in an increased level of DBSs. **A**. Venn diagram shows overlap among DNA DSB break cluster sites in Jurkat WT, KO1, and KO2 cells. The total numbers of break cluster sites in each sample are shown. **B.** Mean coverage of DSBs at merged break cluster sites (n = 33 304) are analyzed. **C.** The DSB coverage at the merged break cluster sites and randomly shuffled control sites (n = 33 304) shows that the WT exhibited a high level of DSBs (compared to the randomly shuffled regions), and the DSBs were further increased upon loss of DHX36 helicase (** *p* < 0.01, *** *p* < 0.001, **** *p* ∼ 0, Wilcoxon signed-rank test). Box denotes 25th and 75th-percentiles, the middle bar shows the median, and whiskers span from 5% to 95%. **D.** Heatmaps demonstrate DSBs coverage at the break cluster sites ordered by the DSB coverage of the WT. RPKM – reads per million per kilobases. **E.** Break cluster sites (n = 33 304) mapped in WT and KO cells are preferentially enriched at TSSs (number of peaks assigned to each feature are shown; P – promoter, GB – gene body, IG - intergenic). Genome annotations definitions (TSS – transcription start site, TTS – transcription termination site).

### DNA double-strand breaks are enriched at G4 genomic regions in DHX36 KO cells

To investigate whether the break cluster sites observed in DHX36 KO cells were enriched in regions containing G4 structures, we used two complementary approaches. First, we identified G4 consensus sites across six different cell lines (NHEK, HaCaT, U2OS, K562, HeLa, and K562) using ChIP-seq and CUT&Tag data generated with the structure-specific antibody, BG4 (6, 8, 13, 20, 65, 66). This analysis identified 4 880 G4 consensus sites present in at least five out of six cell lines, with 88% of these sites overlapping with the break cluster sites mapped in Jurkat cells (**Figure 2A**). The loss of DHX36 helicase led to a significant enrichment of DSBs at G4 sites when compared to WT and randomly shuffled control (**** *p* ∼ 0, ns – not significant, Wilcoxon signed-rank test, **Figure 2B**). A heatmap representation further highlighted this enrichment of DSBs at G4 sites, supporting the notion that G4 structures contribute to genomic instability under normal conditions and are especially prone to DSBs when unresolved in DHX36-depleted cells (**Figure 2C**). Notably, G4 consensus sites were preferentially located at transcription start sites (**Figure 2D**), mirroring the distribution of break cluster sites in DHX36 KO cells (**Figure 1E**). This result is consistent with previous findings that G4 structures are enriched at the active promoter regions and influence gene expression. To further investigate the nature of DSBs at the G4 structures, we used the non-B database (58) to extract the canonical G-quadruplex-forming sequences (G_≥3_ N_1-7_ G_≥3_ N_1-7_ G_≥3_ N_1-7_ G_≥3_ N_1-7_) located within break cluster sites. Comparison of DSB coverage revealed that break cluster sites containing G4 motifs exhibited significantly higher DNA fragility than those lacking canonical G4 motifs (**\*\*\*\*** *p* ∼ 0, Wilcoxon signed-rank test, **Figure 2E**). Among sites with G4 motifs, we observed a bias towards DNA breakage at the C-rich strand that does not form the G4 structure (an example shown in **Supplementary Figure S4A**). This strand-specific enrichment was statistically significant (*** *p* < 0.001, **** *p* ∼ 0, Kruskal-Wallis, post-hoc Dunn’s test, **Figure 2F**) and further potentiated by DHX36 loss. These findings indicate that cycling cells naturally maintain low basal levels of DSBs at G4 structures, a process tightly regulated by DHX36 helicase. The increased DNA fragility at G4 structures in DHX36 KO cells underscores its critical role as a genomic guardian, preventing naturally occurring DSBs at these sites.

**Figure 2.**
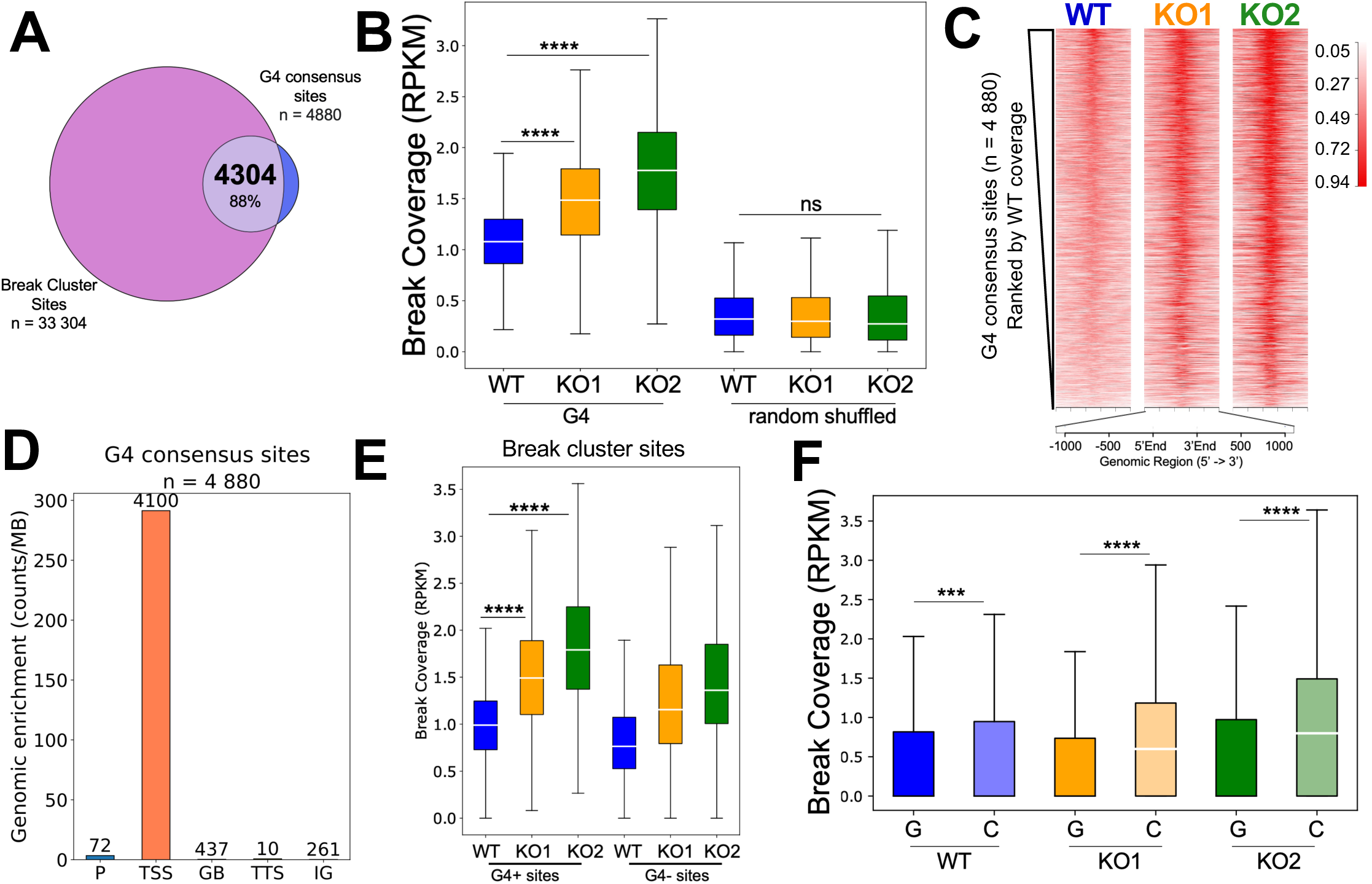
DSBs enrichment at the G4 consensus sites upon loss of DHX36 helicase. G4 consensus sites (n = 4 880) are shared across different cell types and were detected *in vivo* by the structure-specific antibody BG4. **A.** Venn diagram shows overlap between identified break cluster sites and G4 consensus sites. **B**. DHX36 KO cells have significantly more DSBs mapped at G4 consensus sites, compared to the WT (**** *p* ∼ 0, Wilcoxon signed-rank test). Break coverage at the randomly shuffled control G4 regions (n = 4 880) was not statistically significant between the WT and KO cells (ns, Wilcoxon signed-rank test). **C.** Heatmaps demonstrate DSB coverage over G4 consensus sites ordered by the DSB coverage of the WT. **D.** G4 consensus sites are preferentially located at TSSs (number of peaks assigned to each feature are shown; P – promoter, GB – gene body, IG - intergenic). **E.** Boxplot shows quantification of break coverage at break cluster sites that contain G4 motif (G4+, n = 6 834) and the rest of the sites (G4-, n = 26 470) (**** *p* ∼ 0, Wilcoxon signed-rand test). **F.** Mapped G4 motifs present in break cluster sites (n = 9013) showed significant enrichment in DSBs at the C-rich strand, compared to the G4 strand, and this enrichment is further exacerbated by DHX36 loss (*** *p* < 0.001, **** *p* ∼ 0, Wilcoxon signed-rank test). RPKM – reads per kilobases per million. Box denotes 25^th^ and 75^th^-percentiles, middle bar shows the median, and whiskers span from 5% to 95%.

### G4 structures promote genome instability at gene regulatory regions in a function-dependent manner

Enrichment of DSBs and G4 consensus sites at transcription start sites prompted us to investigate genome instability in gene regulatory regions associated with active transcription. Highly expressed genes (n = 2 548) displayed enrichment of DSBs at TSSs in wild type cells, consistent with previous reports (53, 56). Loss of DHX36 helicase significantly amplified DSB enrichment at these TSSs (**\*\*\*\*** *p* ∼ 0, Wilcoxon signed-rank test, **Supplementary Figure S5A &B**). In contrast, low expressed genes (n = 2 543) did not show a significant DSB enrichment, implying a dependence on active transcription (67). One of the mechanisms of gene expression regulation is RNA polymerase (Pol) II promoter-proximal pausing, and it has been shown that genes where RNA Pol II pauses correlate with the presence of more than one G4 motif (68, 69). Accordingly, we observed a significant increase in DNA breaks at RNA polymerase II pausing sites (n = 7 941) in DHX36 KO cells compared to wild type or randomly shuffled pausing sites (**\*\*\*\*** *p* ∼ 0, ns – not significant, Wilcoxon signed-rank test **Supplementary Figure S5C &D**). These findings align with the structural predisposition of G4 motifs to form energetically stable secondary structures, contributing to genomic instability. Next, we examined histone marks, H3K27ac and H3K4me3, which are associated with active enhancers and promoters and are known to colocalize with G4 structures (70, 71). Strong H3K27ac sites showed significant DSB enrichment compared to weaker sites, a pattern exacerbated by DHX36 loss (**\*\*\*\*** *p* ∼ 0, Wilcoxon signed-rank test, **Supplementary Figure S5E & F**). We observed a similar trend for H3K4me3 (**\*\*\*\*** *p* ∼ 0, Wilcoxon signed-rank test, **Supplementary Figure S5G & H**) and CTCF-binding sites (**\*\*\*\*** *p* ∼ 0, Wilcoxon signed-rank test, **Supplementary Figure S5I & J**), which regulate chromatin architecture and transcriptional insulation (72).

To determine whether G4 motifs drive DSB enrichment at these regulatory regions, we categorized TSSs, H3K27ac sites, H3K4me3 sites, and CTCF-binding sites into G4+ (containing G4 motifs), and G4− (lacking G4 motifs) groups. G4+ sites showed significantly higher DSB levels in DHX36 KO cells than WT cells (**\*\*\*\*** *p* ∼ 0, Wilcoxon signed-rank test, **Figure 3A-D**). This trend persisted across TSSs, H3K27ac, H3K4me3, and CTCF-binding sites, with G4+ sites consistently exhibiting higher DNA fragility than G4− sites (**Figure 3B-D**). These results establish G4 structures as key drivers of genome instability, an effect that becomes more pronounced when DHX36 helicase is lost.

**Figure 3.**
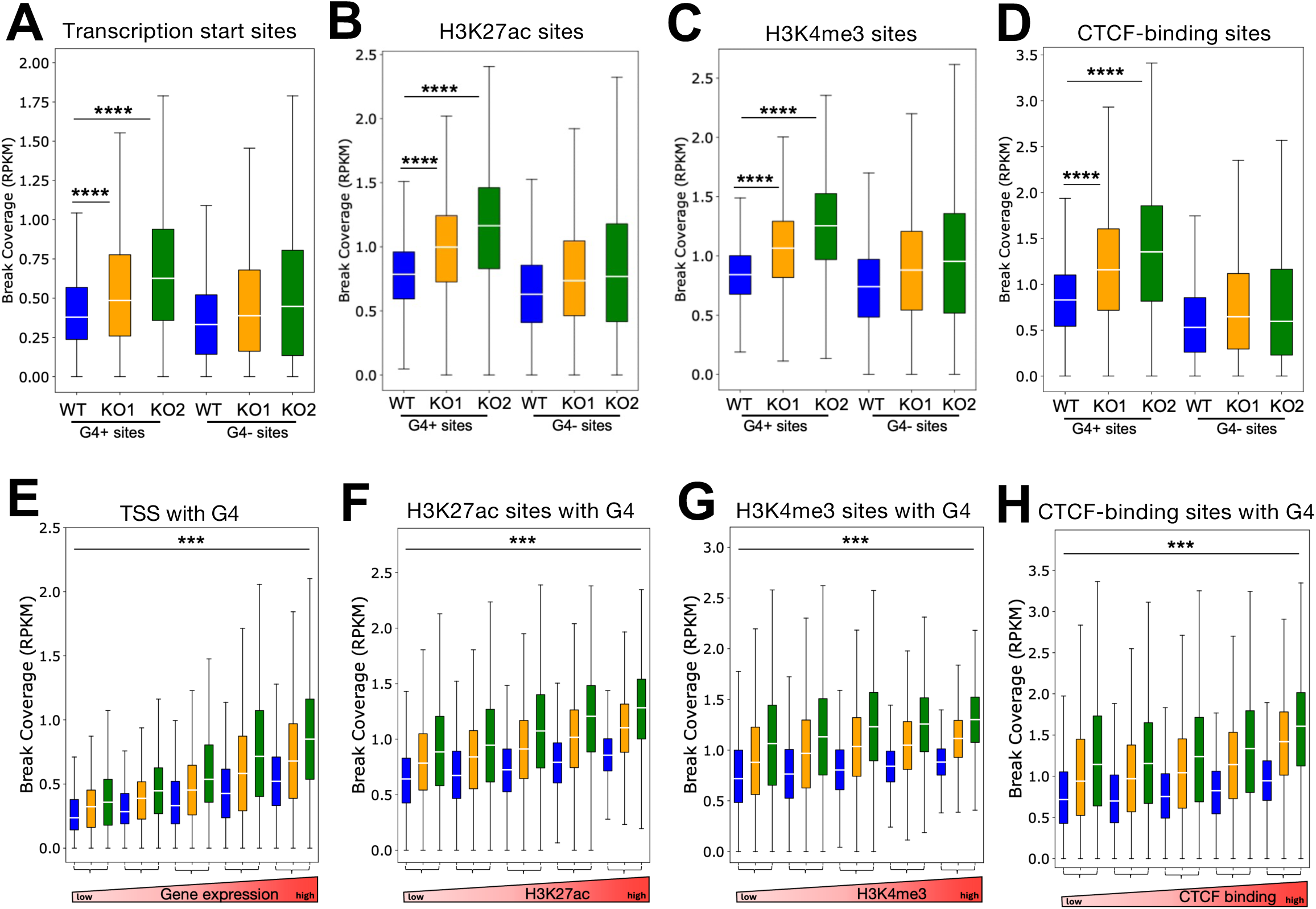
Genome instability at G4-containing sites upon loss of DHX36 is dependent on functional activity of the sites. Boxplot shows quantification of break coverage at (**A**) transcription start sites +/-250 bp that contain a canonical G4 motif (G4+, n = 8 628) and the remaining transcription start sites (G4-, n = 14 282); (**B)** H3K27ac sites that contain a canonical G4 motif (G4+, n = 15 352) and the rest of the sites (G4-, n = 44 121); (**C)** H3K4me3 sites that contain a canonical G4 motif (G4+, n = 12 216) and the rest of the sites (G4-, n = 16 236); (**D**) CTCF-binding sites that contain a canonical G4 motif (G4+, n = 7 106) and the rest of the sites (G4-, n = 61 097) (**** *p* ∼ 0, Wilcoxon signed-rand test). **E.** Break coverage at G4+ TSS sites binned by the gene expression showed a transcription-dependent genome instability. Break coverage at G4+ H3K27ac (**F**), H3K4me3 (**G**), and CTCF-binding (**H**) sites binned by the respective macs2 score shows a correlation between DSB levels and the abundance of histone marks or CTCF binding strength (**\*\*\*** *p* < 0.001, Kruskal-Wallis, post-hoc Dunn’s test). The strongest regulatory elements showed the highest levels of DSBs, particularly in DHX36 KO cells. RPKM – reads per kilobases per million. Box denotes 25^th^ and 75^th^-percentiles, middle bar shows the median, and whiskers span from 5% to 95%.

We further examined whether the regulatory strength of these sites influences DSB levels. At TSS G4+ sites, we observed significant increasing DSBs in accordance with increasing gene expression in the WT cells, with further increases in DHX36 KO cells (**\*\*\*** *p* < 0.001, Kruskal-Wallis, post-hoc Dunn’s test, **Figure 3E**). Similarly, G4+ H3K27ac sites (**Figure 3F**), H3K4me3 sites **(Figure 3G**), and CTCF-binding sites **(Figure 3H**) exhibited a positive correlation between DSB levels and the abundance of histone marks or CTCF binding strength (**\*\*\*** *p* < 0.001, Kruskal-Wallis, post-hoc Dunn’s test). The strongest regulatory elements showed the highest levels of DSBs, particularly in DHX36 KO cells (**Figure 3E-H**). Our findings demonstrate that G4 motifs at key regulatory regions, including H3K27ac, H3K4me3, and CTCF-binding sites, drive genome instability. The presence of G4 structures, coupled with functional activity at these sites, creates hotspots of DNA fragility, and the loss of DHX36 helicase significantly exacerbates this instability.

### Depletion of DHX36 helicase leads to upregulation of genes involved in the immune response

The significant enrichment of DNA double-strand breaks at transcription-related regions, and our previous observation that genes undergoing active transcription are enriched in DSBs at the TSS regions (53, 56) prompted us to explore how the loss of DHX36 impacts transcriptional regulation. We conducted RNA-seq analysis to determine whether DHX36 depletion affects gene expression. The principal component analysis confirmed that the loss of DHX36 accounted for the majority of the variance (81%) in the data (**Supplementary Figure S6A**). We used DESeq2 to identify genes with expression that are significantly altered upon loss of DHX36 (**Supplementary Figure S6B & C**). The correlation between differentially expressed genes in KO1 and KO2 was high (Pearson’s correlation r = 0.75, p ≅ 0, **Figure 4A**). Loss of DHX36 led to significant changes (FDR < 0.05 and abs(log_2_FC) ≥ 1) in gene expression of 1629 and 1603 genes that showed differences in WT vs. KO1 and WT vs. KO2, respectively. The number of significantly upregulated genes in KO1 and KO2 was 686 and 583, respectively (FDR < 0.05 and log_2_FC ≥ 1), and a large overlap between them was observed (n = 423, **Figure 4B**). Gene ontology (GO) analysis of these significantly activated genes revealed enrichment in immune-related processes, including T cell activation, adaptive immune response, leukocyte cell-cell adhesion, and cytokine production (**Supplementary Figure S7A & B**). Conversely, 943 and 1,020 genes were significantly downregulated genes upon loss of DHX36 (FDR < 0.05 and log_2_FC ≤ −1). These genes were enriched in biological processes related to cell junction organization and locomotion (**Supplementary Figure S7C & D**). Interestingly, genes upregulated following the loss of DHX36 demonstrated significant GC enrichment (*** *p* < 0.001, Chi-squared test, **Supplementary Figure S8A & B**). In contrast, downregulated genes showed no significant GC enrichment (ns – not significant, Chi-squared test, **Supplementary Figure S8C & D**). These results suggest that DHX36 helicase plays a role in regulating immune signaling pathways and mRNA transcripts of GC-rich genes in Jurkat cells.

**Figure 4.**
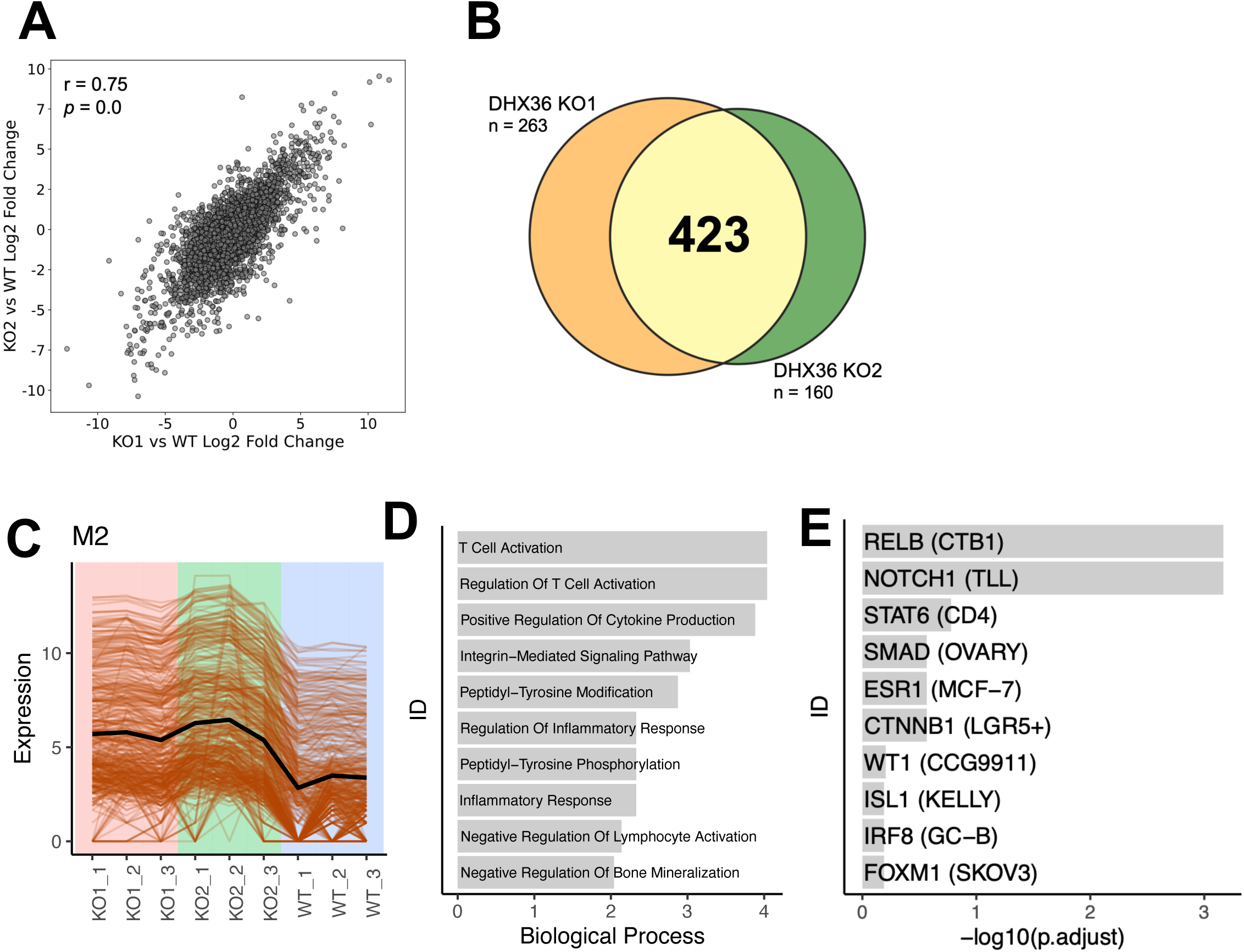
Depletion of DHX36 leads to the activation of genes involved in immune response in T-lymphoblastoid cells. **A.** The gene expression difference between DHX36 KO1 vs WT and DHX36 KO2 vs WT showed a strong correlation (Pearson’s correlation r = 0.75, p ≅ 0). **B**. Significantly upregulated genes in DHX36 KO1 (n = 686 total) and DHX36 KO2 (n = 583 total) show a large overlap, n = 423**. C.** Expression profile shows the 401 genes in Module M2 that are upregulated in the DHX36 KO cells compared to the WT. **D.** Genes in Module M2 show significant enrichment of immune cell activation-related biological processes. **E.** Genes in M2 are enriched for targets of RELB and NOTCH1.

To further de-convolute common biological themes between the two DHX36 KO cell lines, we used the co-expression modules identification tool (CEMiTool) to identify systems-level gene modules of co-varying genes. CEMiTool identified 11 modules. The module with the largest number of genes, M1, included 465 genes with higher expression in the WT samples than in the KO cell lines (**Supplementary Figure S9A**). Enrichment analysis showed that M1 genes were gene targets of T-Cell Acute Lymphocytic Leukemia Protein 1 (TAL1/SCL), a helix-loop-helix transcription factor that is an essential regulator in hematopoietic cells, and Retinoid X receptors (RXR), an important member of the nuclear receptor superfamily required for T-cell differentiation, function, and tissue homing (**Supplementary Figure S9B**). The second-largest module, M2, included 401 genes with higher expression in the DHX36 KO cell lines than the WT (**Figure 4C**). This module was enriched for genes involved in T-cell activation, cytokine production, and inflammatory response (**Figure 4D**). Transcriptional response leading to the activation of immune gene expression is governed by numerous transcription factors that reside downstream of multiple immune signaling pathways. Enrichment analysis of genes in module M2 against CHEA Transcription Factor Targets identified RelB (a member of the NF-κB pathway essential for T-cell development) and NOTCH1 (a transmembrane receptor critical for T-cell fate and function) (**Figure 4E**). These findings indicate that DHX36 depletion upregulates immune-related transcriptional programs, including NF-κB driven pathways, connecting DHX36 helicase to the regulation of canonical and non-canonical NF-κB signaling.

### Loss of DHX36 leads to the activation of NF-κB signaling pathway

The activation of the NF-κB signaling pathway is a common downstream event of diverse pattern recognition receptors (PRRs), driving the transcription of pro-inflammatory cytokines, chemokines, and other inflammatory mediators (73, 74). The NF-κB transcription factor family, including RelA/p65, RelB, p50, p52, and c-Rel, regulates gene expression by binding specific DNA elements (73). In unstimulated cells, NF-κB (p50:RelA/p65) resides in the cytoplasm, and upon phosphorylation and degradation of the inhibitory κB protein, NF-κB translocates into the nucleus to activate gene expression (73, 74). To investigate NF-κB dynamics, protein fractionation experiments were performed to quantitate RelA/p65 levels in the cytoplasm and nucleus of Jurkat cells (**Figure 5A**). DHX36 KO cells showed a significant accumulation of nuclear RelA/p65 relative to WT cells (* *p* < 0.05, *** *p* < 0.001, Student’s t-test, **Figure 5B**), indicating that RelA/p65 drives immune gene activation in this context. Systems-level analysis as described in **Figure 4**, revealed upregulation of NF-κB signaling pathways in DHX36 KO cells, with 76% of NF-κB target genes enriched relative to WT cells (**Figure 5C**). To validate these findings from RNA-seq, we performed RT-qPCR on selected DHX36 KO upregulated genes. We examined known NF-κB regulated immune genes with BG4-detected G4 structures within their proximal promoter regions. DHX36 KO cells significantly upregulated *B2M*, *IL16*, *IL32,* and *TNFSF10* genes compared to WT cells (* *p* < 0.05, ** *p* < 0.01, Student’s t-test, **Figure 5D**). We performed a Luminex assay to measure supernatant-secreted cytokines to determine whether the observed transcriptional signature of pro-inflammatory cytokines in DHX36 KO cells led to their production and secretion. CCL1/I-309 and IL-16, two potent pro-inflammatory ligands, were produced and secreted in higher amounts in DHX36 KO Jurkat cells compared to the WT (**Figure 5E**). Given these observations, we explored whether loss of DHX36 and increased pro-inflammatory cytokine production involve the cGAS-STING immune signaling pathway, which could be influenced by the observed level of DSBs detected in DHX36 KO cells. Western blot and RT-qPCR analyses revealed significantly elevated STING protein (* *p* < 0.05, ** *p* < 0.01 Student’s t-test, **Figure 5F**) and *STING1* mRNA levels (** *p* < 0.01, Student’s t-test, **Figure 5G**), suggesting a role for STING in driving the observed immune-related gene signature.

**Figure 5.**
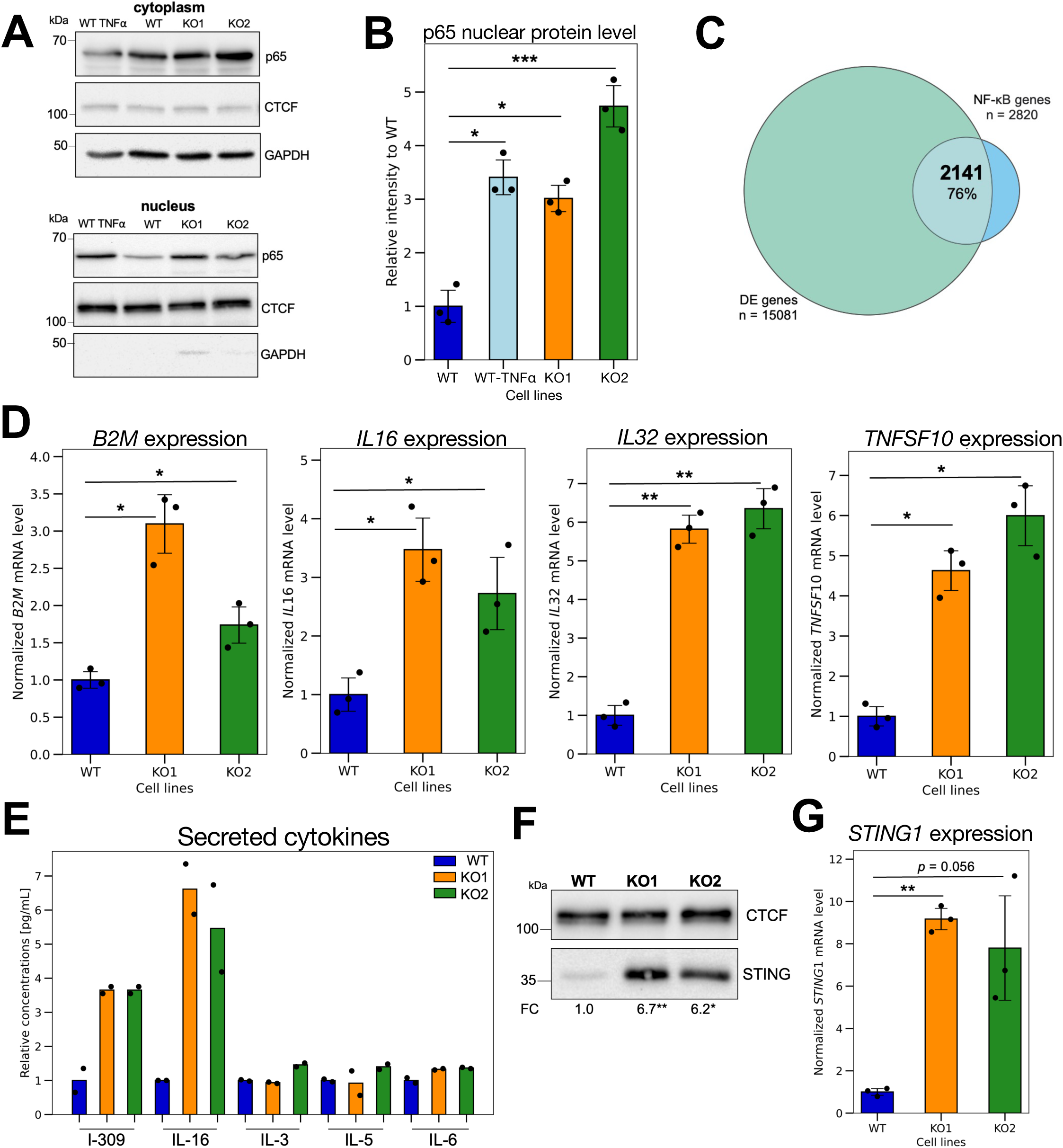
Loss of DHX36 leads to the activation of the NF-κB signaling pathway. **A.** Western blot of p65 presence in the cytoplasm and nucleus shows that p65 is translocated to the nucleus upon loss of DHX36. Treatment of the WT cells with TNFα, known as NF-κB activator, was used as the positive control. **B.** In DHX36 KO cells, the p65 level in the nucleus is significantly higher compared to WT cells (* *p* < 0.05, *** < 0.001, Student’s t-test). **C.** Differentially expressed genes upon DHX36 knock-out are enriched for the NF-κB target genes (n = 2141, 76%). **D.** At the mRNA level, loss of DHX36 leads to a statistically significant upregulation of NF-κB target genes involved in activation of immune response, such as *B2M, IL16, IL32, TNFSF10* (* *p* < 0.05, ** *p* < 0.01, Student’s t-test). **E**. Luminex assay of supernatant-secreted cytokines shows that CCL1/I-309 and IL-16, two potent pro-inflammatory ligands, were produced and secreted in higher amounts in DHX36 KO cells compared to the WT. Two independent biological replicates were shown. **F.** Significantly higher level of STING protein upon loss of DHX36 (FC – fold change, * *p* < 0.05, ** *p* < 0.01, Student’s t-test, n = 3 each); protein level was normalized to the loading control and the WT signal. **G**. At the mRNA level, loss of DHX36 leads to the upregulation of the *STING1* gene as shown by RT-qPCR (** *p* < 0.01, Student’s t-test, n = 3 each).

### Better survival of B-ALL patients correlates with high expression of *DHX36*

DHX36 is essential in many developmental processes (46–48) and is often mutated across different cancer types (75). Its gene expression is frequently elevated in cancers, prompting us to compare DHX36 expression across tumor cohorts in the Genomic Data Commons (**Supplementary Figure S10A**). TARGET-CCSK (Clear Cell Sarcoma of the Kidney) showed the highest median *DHX36* expression, but has a limited number of patients (n = 13). Therefore, we focused on the TARGET-ALL (Acute Lymphoblastic Leukemia) cohort because it has the second-highest *DHX36* median expression and robust sample size (n = 108). We stratified the 108 B-ALL bone marrow patient samples into HIGH and LOW groups based on the *DHX36* expression. Using differential gene expression analysis (accounting for gender and cohort differences), we identified 6 505 genes that were significantly upregulated (FDR < 0.05 and log_2_FC ≥ 1) and 2 814 genes that were significantly downregulated (FDR < 0.05 and log_2_FC ≤ −1) in the DHX36-HIGH group compared to the DHX36-LOW group (**Supplementary Figure S10B**). To identify molecular signatures of these differentially expressed genes, we performed a GSEA analysis, and found that genes upregulated in the DHX36-HIGH group were enriched in processes associated with mitotic spindle and TGFβ signaling. In contrast, genes downregulated in the DHX36-HIGH group were enriched for gene sets associated with response to interferons and DNA repair (**Supplementary Figure S10C**), supporting the notion that DHX36 maintains genome integrity and concurring with our findings in DHX36 KO cells. We also observed the expression of *STING1* to be significantly higher in patients with lower *DHX36* gene expression (** *p* < 0.01, Wilcoxon signed-rank test, **Figure 6A**), consistent with our observation in DHX36 KO cells (**Figure 5G**). We next investigated whether changes in *DHX36* gene expression affected survival outcomes for B-ALL patients. We found that lower *DHX36* expression is associated with a worse prognosis in B-ALL patients (*** *p* < 0.001, log-rank test, **Figure 6B**). To further understand the underlying mechanism for this difference in survival, we de-convoluted the bulk gene expression data using xCell, a cell-type enrichment analysis. The *HIGH* DHX36 expression group showed significantly higher enrichment for natural killer T (NKT) cells, which is associated with a better prognosis (76). This observation of significantly a lower NKT cell score in the DHX36-LOW group (*** *p* < 0.001, Wilcoxon signed-rank test, **Figure 6C**) is consistent with the results of our co-expression module analysis in the DHX36 KO cells (**Figure 4D**). The M2 module enrichment analysis shows enrichment of genes associated with decreased NKT cell number in DHX36 KO cells. These findings show that the lower level of *DHX36* is strongly associated with DNA damage and higher expression of *STING1* and with a less favorable prognosis in the TARGET pediatric B-ALL tumor samples, further highlighting the essential role of DHX36 in genome stability.

**Figure 6.**
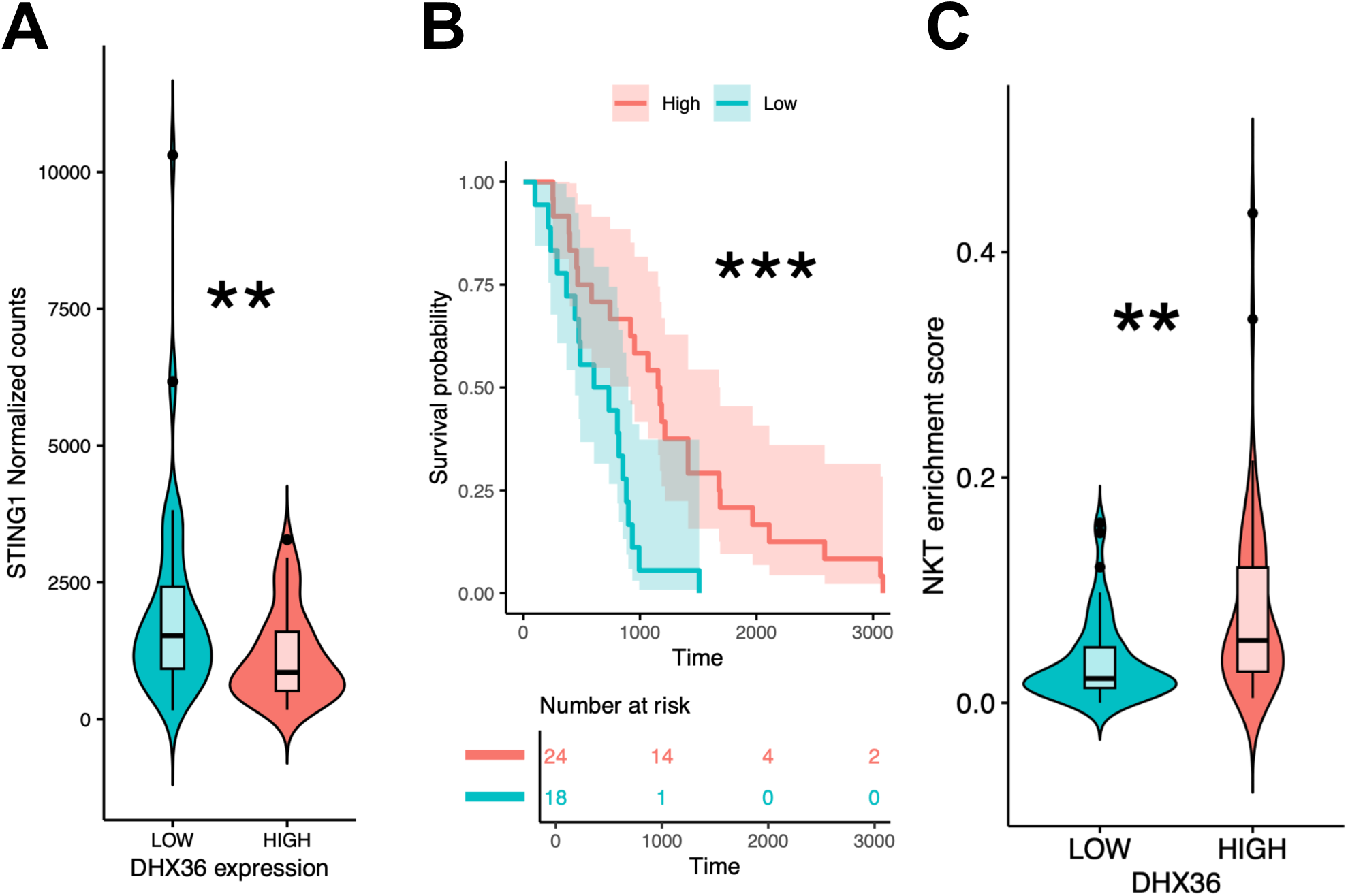
Better outcomes for B-ALL patients with high expression of *DHX36*. **A.** Low expression of *DHX36* correlates with high expression of *STING1* in B-ALL patients (** *p* < 0.01, Wilcoxon signed-rank test). **B.** Lower *DHX36* expression leads to worse survival in pediatric B-ALL based on a KM test (*** *p* < 0.001, log-rank test). This difference remains significant in a Cox-proportional hazards model after accounting for age at diagnosis and gender, which are known to affect survival in B-ALL. **C.** Higher *DHX36* expression correlates with higher natural killer T cells (NKT) enrichment (** *p* < 0.01, Wilcoxon signed-rank test).

## DISCUSSION

We demonstrated that the loss of DHX36 helicase significantly contributes to DSBs and genome instability, resulting in the activation of immune genes in T-cells. The data strongly indicates that unresolved G4 structures are a source of endogenous DNA breaks in active genes enriched for CTCF, H3K4me3, and H3K27ac modifications associated with transcription. Most (88%) G4 conserved sites (present in five out of the six cell lines) are identified as break cluster sites in DHX36 KO cells, which agrees with previous reports linking G4 and genome instability (42). Moreover, upon DHX36 depletion, we have shown strong T-cell activation at the transcriptional level, driven by the NF-κB signaling pathway and the production of pro-inflammatory cytokines. Additionally, we observed that pediatric B-ALL patients with low expression of DHX36 have worse outcomes, highlighting the importance of this helicase in maintaining genome stability.

Using DSB genome-wide break mapping assays, our results not only highlight the importance and extent by which DHX36 protects the cell from DSBs near G4 structures but also allow for a better understanding of why this regulation is so important. Mizumoto *et al*. (42) first demonstrated that the knockdown of DHX36 in IMR-90 cell lines resulted in the accumulation of DNA damage detected by immunofluorescence of γH2AX, which colocalizes with BG4 staining. However, such approaches lack high resolution; therefore, we applied single-nucleotide DSBs mapping, which provided molecular insight into DHX36/G4-genome instability, with the ability to investigate not only the genomic location of DNA breakage but the extent of those events and their consequences at the transcriptional level. We showed enrichment of DNA breaks in DHX36 KO cells colocalized with the G4 site, p65 occupancy, and RNA Pol II pausing sites at the TSSs of *B2M* and *NOTCH1* genes, and these chromatin-associations correspond with significant upregulation of mRNA expression of those genes (**Figure 5D and Supplementary Figure S11A and B).** Our work highlights the importance of DHX36 as a G4 resolvase that not only protects from DSBs but also regulates cellular gene expression involved in the activation of innate immune signaling pathways. Because of the significant enrichment of NF-κB regulated genes following the loss of DHX36 expression, we postulate that one of the important roles of DHX36 is to control inappropriate innate immune signaling mediated by NF-κB activation. Since the loss of DHX36 is known to increase G4 structures (41, 42), perhaps NF-κB serves a unique role in bypassing RNA Pol II pausing of gene promoters rich in G4 structures. In support of this hypothesis, NF-κB aids in the release of RNA Pol II pausing (77, 78) and p65 has recently been shown to directly bind G4 structures to control glycolytic gene expression in mitochondria (79).

DNA G4s can both positively and negatively regulate transcription. Our observations that the loss of DHX36 results in both an enrichment in DSBs at TSSs and an increase in mRNA encoding transcripts from these genes suggest that there are a number of genes in which increased DSBs promote active transcription. We speculate that DNA breaks enriched at transcription start sites could be linked to the role of DNA topoisomerase II (TOP2) in resolving transcription-associated supercoiling. Given that TOP2 can fail to re-ligate DNA strands endogenously, leading to trapping TOP2 cleavage complexes (53, 80), our findings align with previous work by Olivieri *et al.* (81). Using an unbiased genetic screen, they reported that stabilization of G4 by small ligands can trap TOP2 on DNA and knocking down DHX36 sensitize the cells to TOP2 poisons. These observations support a mechanism where transcription-associated genome instability upon loss of DHX36 results from trapping TOP2 at G4 sites. Furthermore, DSBs at and near transcription start sites have been shown to link to the release of RNA Pol II promoter-proximal pausing, leading to increased gene expression (13, 56, 82, 83). The enrichment of DSBs at the RNA Pol II promoter-proximal pausing sites in the WT cells and its increase in the DHX36 KO cells shown in **Supplementary Figures S5C and S5D** support the involvement of DSBs in pausing release. Furthermore, TOP2 activity is elevated in active chromatin compartments and associated with CTCF and histone modifications observed during transcription (H3K4me3 and H2K27ac) (84, 85). CTCF physically interacts with TOP2 and recruits to the DNA to relieve torsional stress during transcription (86, 87). Our recent study on DSBs at CTCF binding sites suggests that G4s act as a recognition sequence for TOP2 binding and cleavage at CTCF binding sites (52). These observations support the mechanisms where transcription-associated genome instability upon loss of DHX36 results from trapping TOP2 at G4 sites to initiate DSBs.

Upon loss of DHX36, we observed overlap between identified break cluster sites and G4 consensus sites (**Figure 2A**), which agrees with previous reports linking G4 and genome instability (42). The advantage of employing a sequencing strategy to investigate genome instability revealed that C-rich strands within G4 motifs have a higher predisposition to breakage (**Figure 2F & Supplementary Figure S4A**). This observation is intriguing and suggests that G4s are unfavorable for DNA breaks, possibly due to the structure itself and/or the binding of protective protein factors. The DNA breaks occurring at the flanking sequences of the G4s can promote the generation of DNA fragments ready to form G4s; as shown in Byrd et al. that oxidative stress-induced DNA damage promoted G4-containing fragments excised from the genome and released into cytoplasm which can be detected by the structure-specific BG4 antibody (88). The bias towards DNA breaks at C-rich strands also raises the possibility of the involvement of DNA:RNA hybrids and the stabilization of this structure by the formation of G-loop (89, 90), which negatively correlates with transcription (17, 91). ZNF*672,* which displays the C-rich strand DSB bias (**Supplementary Figure S4A**), is downregulated in gene expression upon loss of DHX36, possibly serving as an example (**Supplementary Figure S4B**). Further studies of the mechanism for the DNA break bias on C-rich strands are warranted.

The DHX36 helicase is able to resolve G4 structures in both DNA and RNA (35–38). Significant changes in gene expression that we observed might result in the role of DHX36 in regulating RNA G4 structures present within mRNA transcript (50). DHX36 is a helicase that acts on both DNA and RNA G4s. Sauer *et al*., in a system-wide manner, analyzed mRNA DHX36 targets (41) and found that loss of DHX36 resulted in the accumulation of translationally incompetent target mRNAs. In addition, the depletion of RNA-G4 unwinding helicases such as DHX9 and DHX36 promotes the translation of RNA-G4 associated upstream reading frames while reducing the translation of coding regions for transcripts that comprise proto-oncogenes, transcription factors, and epigenetics regulators (92). Those reposts provide additional evidence of the multifunctional role of DHX36 helicase, where its cytoplasmic and genomic functions should be addressed.

DHX36 has been implicated in the innate immune response through different processes. DHX36 was found to localize in stress granules (SGs) and enhance RIG-I signaling in a stimulus-dependent manner (93). Additionally, DHX36 deficient cells showed defects in IFN production and higher susceptibility to RNA virus infection (94). Zhang *et al*. demonstrated that in murine dendritic cells, DHX36 was found in a complex with other helicases, DDX1 and DDHX21, and acts as a cytoplasmic sensor or viral double-stranded RNA (dsRNA) that uses the TRIF pathway to activate type I IFN responses. Additionally, presented data align with more recent reports from Miglietta *et al*. in which G4 ligands, pyridostatin, and PhenDC at non-cytotoxic dosages showed induction of micronuclei and activation of the cGAS-STING signaling pathway leading to the innate immune gene activation in MCF-7 breast cancer cells (95). This provides strong evidence that manipulation of G4s stability either by small ligands that bind and stabilize G4 or by depletion of DHX36 helicase leads to the cellular activation of an innate immune response.

Despite the presence of sustained accumulation of DSBs, DHX36 KO Jurkat cells adapt, survive, and differentially regulated gene expression, including the upregulation of genes involved in T cell activation and proliferation. Gene set enrichment analysis demonstrates a significant number of the genes upregulated following the loss of DHX36 were well-known NF-κB regulated transcription targets. Not only did DHX36 KO cells display increased accumulation of nuclear p65 component of NF-κB, but cells are functionally activated and secrete cytokines into the growth medium. In addition to driving T cell activation, NF-κB transcriptional programs are known to upregulate gene products that block apoptosis (96). For these reasons, we postulate that as a consequence of DHX36 loss, Jurkat cells upregulate NF-κB as a mechanism to overcome cell death programs mediated by DSBs and proliferation. Thus, additional experiments are needed to determine whether the loss of DHX36 upregulates NF-κB pro-survival function as a compensatory mechanism to avoid cell death.

In conclusion, we provide evidence that DHX36 loss leads to an increased level of DSBs genome-wide and that unresolved DNA G4 structures are prone to DNA breakage at regulatory regions. Loss of DHX36 leads to global changes in gene expression in T cells, which strongly activates the innate immune response through the NF-κB signaling pathway. The results here highlighted DHX36/G4s-driven genome instability and the possibility of targeting DHX36 in cancer cells to stimulate antitumor immunity.

## DATA AVAILABILITY

DSB mapping and RNA-seq datasets generated in this study can be accessed at Sequence Read Archive (SRA): PRJNA795482 for Jurkat WT cells (52) and PRJNA1190447 for DHX36 KO1 and KO2 cells.

## AUTHOR CONTRIBUTIONS

A.R.B., J.P.V., P.J.S., A.R., M.W.M., and Y.-H.W. designed the study. A.R.B. and P.-C.H. performed experiments. A.R.B., A.R., and M.W.M. performed data analysis. A.R.B., A.R., M.W.M., and Y.-H.W. drafted the paper. All authors provided critical review and feedback on the final submission.

