## Supplementary File for "Loss of DHX36/G4R1, a G4 resolvase, drives genome instability and regulates innate immune gene expression in cancer cells"

### Table S1

**Table S1.** Primers for RT-qPCR used in this study.

| Gene target | Forward primer 5’ → 3’ | Reverse primer 5’ → 3’ |
|---|---|---|
| *DHX36* | CCCACCATCAAATGAGGCAGTG | TGTGGCTCAACGGGTAATCGTG |
| *GAPDH* | ACATCGCTCAGACACCATG | TGTAGTTGAGGTCAATGAAGGG |
| *B2M* | GTGCTCGCGCTACTCTCTCT | TTCAATGTCGGATGGATGAA |
| *IL16* | GTCCGTTATCTCCCTGCTGA | AGACCAGCACCTTCCTCCTT |
| *IL32* | TTCAAAGAGGGCTACCTGGA | GGCACCGTAATCCATCTCTT |
| *TNFSF10* | GACAGACCTGCGTGCTGAT | AAGAAACAAGCAATGCCACTT |
| *NOTCH1* | CCAGGAAACAACTGCAAGAA | ACTCGTCCACATCCTCGGTA |

### Table S2

**Table S2.** Sequencing and alignment statistics for 150-bp paired-end Illumina sequencing libraries prepared from Jurkat DHX36 KO1 and DHX36 KO2 cells.

| Cell type | Biological replicates | # Sequenced read pairs[a] | % Alignment rate[b] | # Proper read pairs[c] | % Duplication[d] | # Mapped DSBs[e] | % Mapped DSBs[f] |
|---|---|---|---|---|---|---|---|
| Jurkat DHX36 KO1 | N1 | 24005137 | 89.22 | 20142904 | 22.94 | 14341071 | 59.74 |
| | N2 | 26779482 | 95.27 | 24330205 | 27.25 | 16494690 | 61.59 |
| | N3 | 22444486 | 92.73 | 19437867 | 24.81 | 13333879 | 59.41 |
| | N4 | 27246854 | 93.16 | 23969492 | 20.50 | 17665058 | 64.83 |
| Jurkat DHX36 KO2 | N1 | 17799909 | 92.10 | 15287516 | 40.64 | 8206522 | 46.10 |
| | N2 | 18027974 | 93.09 | 15446217 | 45.37 | 7602851 | 42.17 |
| | N3 | 22660606 | 92.24 | 19488693 | 21.64 | 13875570 | 61.23 |
| | N4 | 24125802 | 91.53 | 20910505 | 21.27 | 15034011 | 62.32 |

[a] Number of raw paired read1-read2s following Illumina paired-end sequencing and quality filtering.
[b] Percentage of reads that had at least one alignment to the hg38 genome assembly as processed and reported by bowtie2 alignment program.
[c] Following the quality control removal of all unmapped, non-primary, supplementary and low-quality reads, the remaining number of paired read1-read2s are indicated.
[d] PCR duplicates are marked and removed meaningfully using read1 and read2 alignment, and "% Duplication" is based on original "# Sequenced read pairs".
[e] For each non-duplicated pair, only read1 is kept, and the 5' most nucleotide of read1 defines the DNA break position.
[f] "% Mapped DSBs" was calculated by dividing "# Mapped DSBs" by "# Sequenced read pairs".

**Table S3.** Publicly available datasets analyzed in this study.

| Species / Cell Line | Dataset Type | Accession(s) | References |
|---|---|---|---|
| *H. sapiens* / NHEK | BG4 ChIP-seq | GSE76688 | Hansel-Hertsch *et al*., Nat Genet 2016 |
| *H. sapiens* / U2OS | BG4 CUT&Tag | GSE181373 | Hui *et al*., Sci Rep 2021 |
| *H. sapiens* / K562 | BG4 ChIP-seq | GSE107690 | Mao *et al.,* Nat Struct Mol Biol 2018 |
| *H. sapiens* / HaCaT | BG4 ChIP-seq | GSE99205 | Hansel-Hertsch *et al*., Nat Protoc 2018 |
| *H. sapiens* / NSC | BG4 ChIP-seq | GSE161531 | Zyner *et al.,* Nat Comm 2022 |
| *H. sapiens* / HeLa | BG4 CUT&Tag | GSE178668 | Li *et al.,* Genome Res 2021 |
| *H. sapiens* / Jurkat | CTCF ChIP-seq | GSE68978 | Hnisz *et al.,* Science 2016 |
| *H. sapiens* / Jurkat | H3K27ac ChIP-seq | GSE68978 | Hnisz *et al.,* Science 2016 |
| *H. sapiens* / Jurkat | H3K4me3 ChIP-seq | GSE23080 | |
| *H. sapiens* / Jurkat | PRO-seq | GSE117832 | Chu *et al.,* Nat Genet 2018 |
| *H. sapiens* / HeLa | RNA Pol II pausing sites | | Szlachta *et al.,* Genome Biol 2018 |
| *H. sapiens* / K562 | p65 ChIP-seq | GSE197704 | |
| *H. sapiens* / Jurkat | DSB mapping/sequencing | PRJNA795482 | Raimer Young *et al.,* NAR 2024 |
| *H. sapiens* / Jurkat | RNA-seq | | |

### Figure S1

| KO1, r | N1 | N2 | N3 | N4 |
|---|---|---|---|---|
| N1 | | 0.980 | 0.965 | 0.989 |
| N2 | 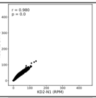 | | 0.922 | 0.964 |
| N3 | 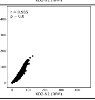 | 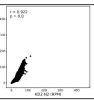 | | 0.982 |
| N4 | 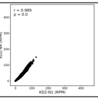 | 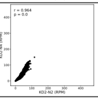 | 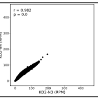 | |

| KO2, r | N1 | N2 | N3 | N4 |
|---|---|---|---|---|
| N1 | | 0.981 | 0.979 | 0.975 |
| N2 | 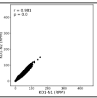 | | 0.979 | 0.974 |
| N3 | 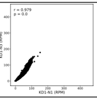 | 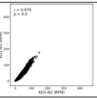 | | 0.995 |
| N4 | 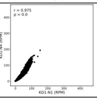 | 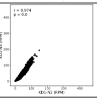 | 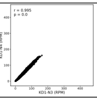 | |

**Figure S1. Reproducibility of genome-wide DSB mapping/sequencing for Jurkat DHX36 KO1 and KO2 cell lines**. Scatter plots of genome-wide DSB mapping/sequencing reads from biological replicates (Table S2) for Jurkat KO1 (left panel) and KO2 (right panel) cells show a strong correlation (KO1 – Pearson's correlation r = 0.922-0.989, p ≅ 0 and KO2 – Pearson's correlation r = 0.974-0.995, p ≅ 0). Read-normalized coverage for each preparation was calculated for 100kb genome-wide, non-overlapping windows (n = 30 895), and Pearson's correlation was calculated.

### Figure S2

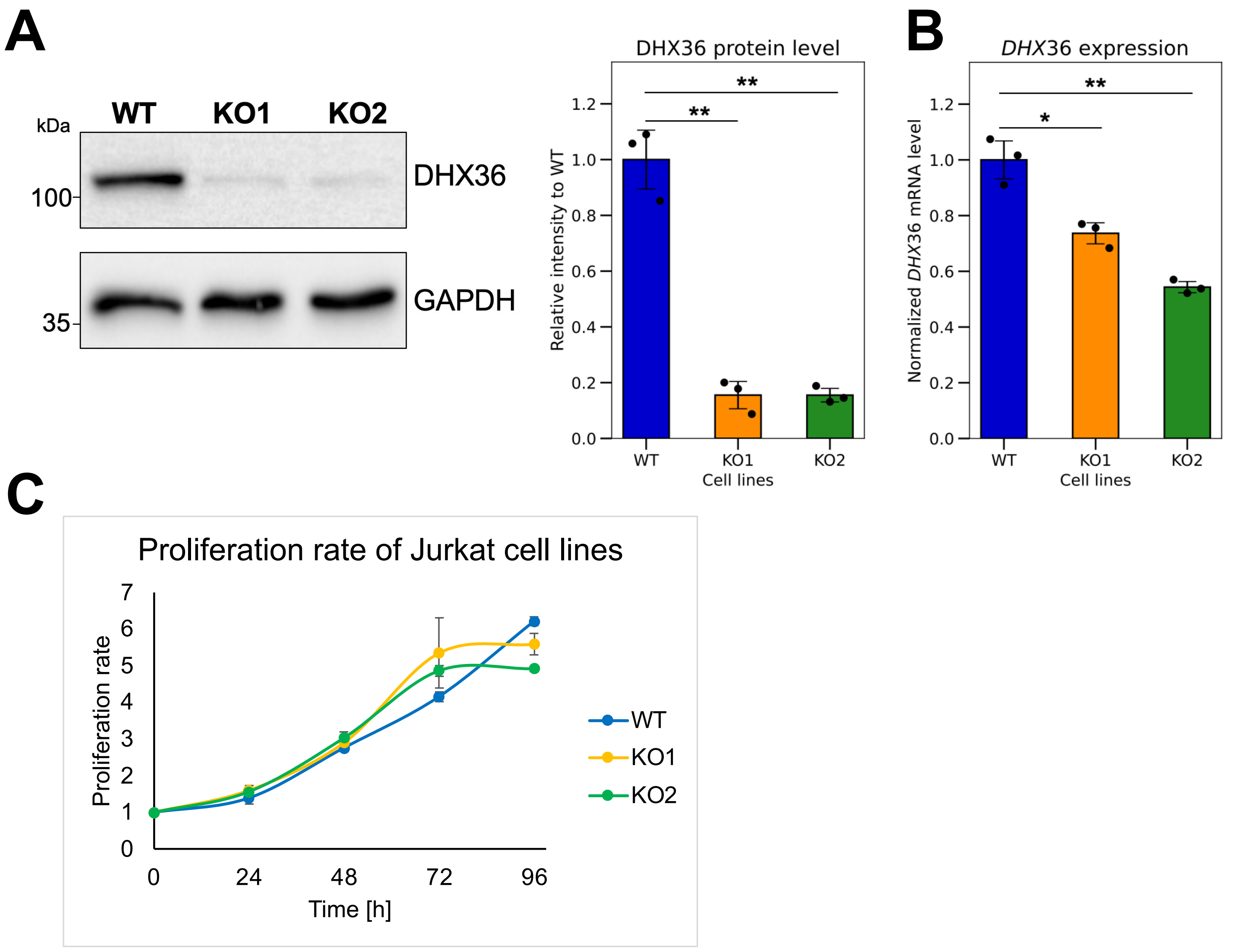


**Figure S2**. **Loss of DHX36 helicase at protein and mRNA level does not affect cell viability**. **A.** Western blot of DHX36 helicase in WT, KO1, and KO2 cell lines. Relative density in each cell line indicates significant knockout of DHX36 at the protein level (n = 3 for each cell line, ** indicates $p < 0.01$, Student's *t* test for pairwise comparisons). N-terminal DHX36 antibody was used. Intensity signal was normalized to the loading control and DHX36 signal from the WT. **B.** At the transcript level, *DHX36* helicase shows significantly lower transcripts compared to WT (n = 3 for each cell line, * indicates $p < 0.05$, ** indicates $p < 0.01$, Student's t-test for pairwise comparisons). **C.** No differences in cell growth were observed between WT, KO1, and KO2 (n = 3 replicates for each) measured by the MTS assay.

### Figure S3

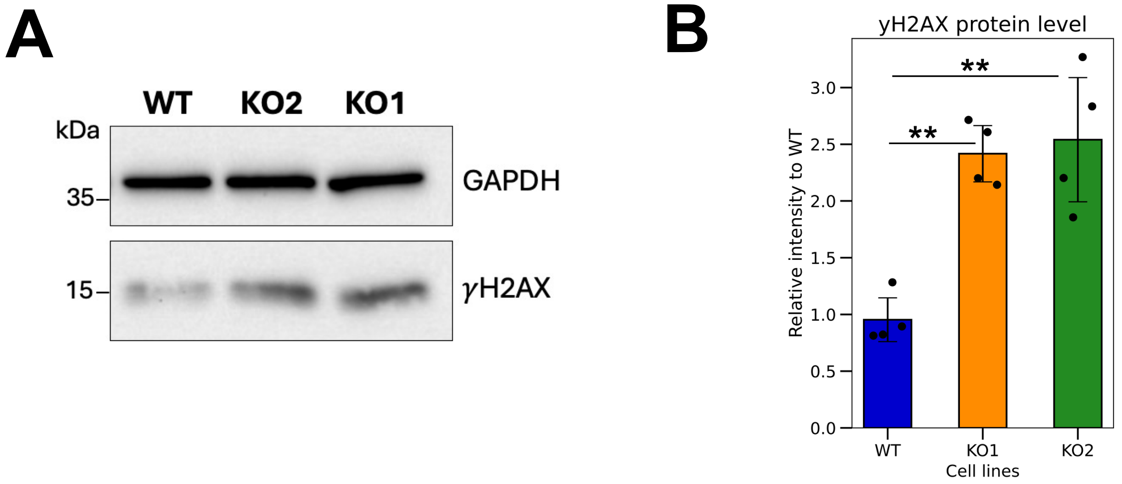


**Figure S3. Significantly higher level of DNA damage response upon loss of DHX36. A.** Western blot of γH2AX, a marker of DNA damage, in Jurkat WT and DHX36 KO cells. **B**. Relative density in each cell line indicates significantly increased γH2AX in KO cells (n = 4 for each cell line, ** indicates $p < 0.01$, Student's t-test for pairwise comparisons). Intensity signals were normalized to their own loading control and γH2AX signal from the WT.

### Figure S4

A

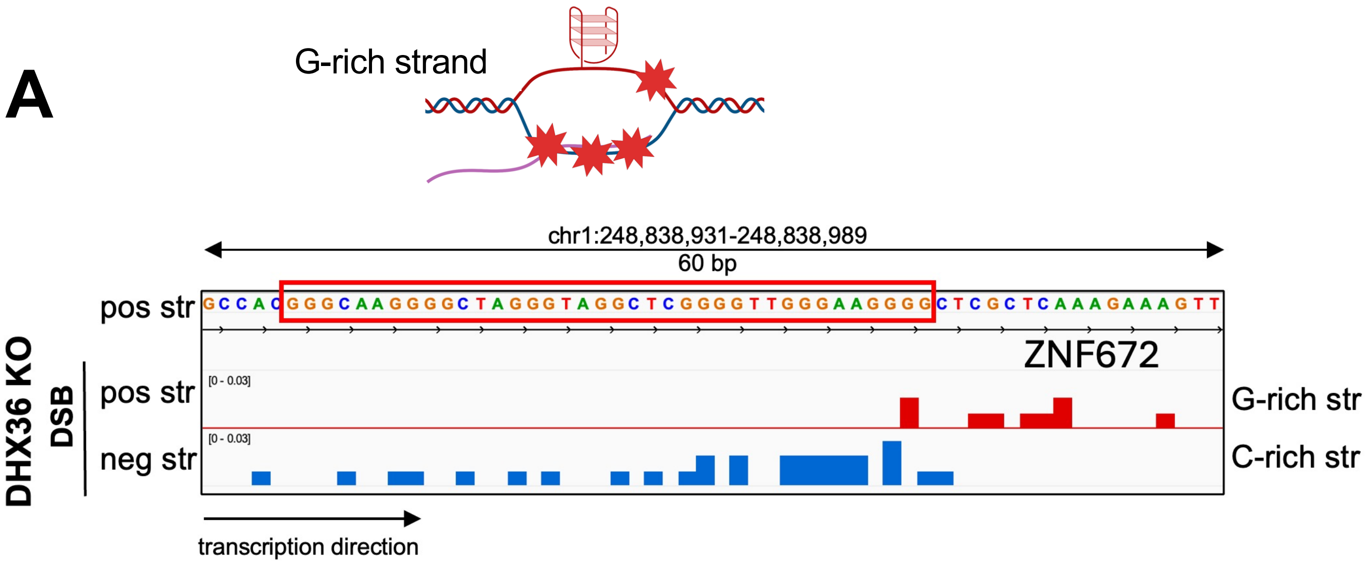


B

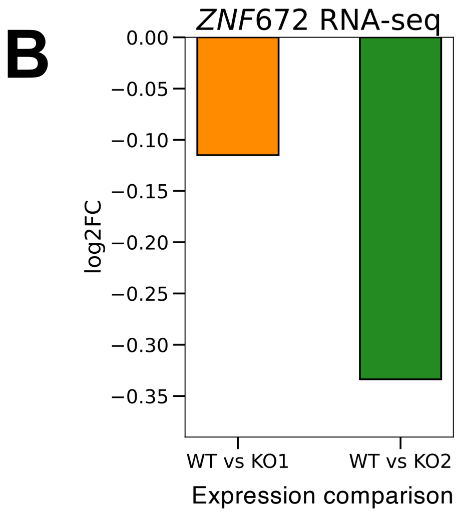


**Figure S4. Preference for DSBs at C-rich strand at G4-containing regions. A.** Example of DNA breaks at G4 region with more breaks at C-rich strand at *ZNF672* gene**. B.** Differential gene expression analysis showed downregulation of *ZNF672* in DHX36 KO cells compared to WT (FC – fold change).

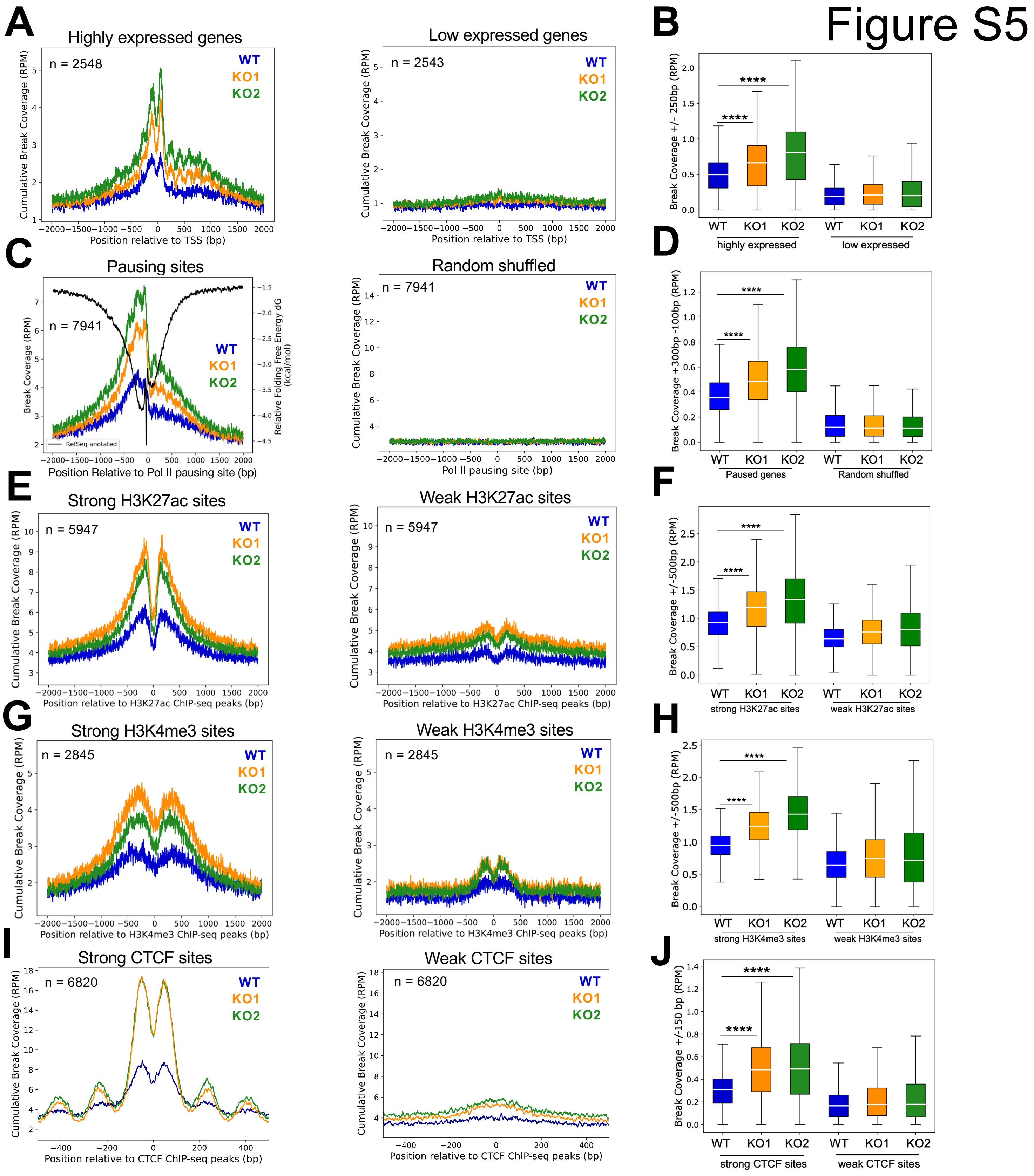


**Figure S5. Enrichment of DSBs at transcription regulated regions. A.** Cumulative, read normalized, single nucleotide resolution profile of DSBs at highly expressed genes (top 10%, n = 2548, left) and at least expressed genes (bottom 10%, n = 2543, right) in WT and DHX36 KO cells. Expression profile is based on the RNA-seq data from WT (n = 3). **B.** Boxplot shows quantification of break coverage at ± 250 bp of TSSs of highly expressed gens and least expressed genes (**** $p$ ~ 0, Wilcoxon signed-rand test). **C.** Cumulative, read normalized, single nucleotide resolution profile of DSBs at RNA Pol II pausing sites (n = 7941, left) in WT and DHX36 KO cells and DNA secondary structure free energy folding (ΔG, kcal/mol)**.** Cumulative DSB coverage at randomly shuffled RNA Pol II pausing sites (n = 7941, right). **D.** Boxplot shows significant enrichment of DSBs at RNA Pol II paused sites upon loss of DHX36 compared to WT (**** $p$ ~ 0, Wilcoxon signed-rand test). For boxplot break coverage regions spanning +300bp and -100bp from RNA Pol II paused site summits were used. **E.** Cumulative, read normalized, single nucleotide resolution profile of DSBs at strong H3K27ac sites (top 10%, n = 5947, left) and at weak H3K27ac sites (bottom 10%, n = 5947, right) in WT and DHX36 KO cells. **F.** Boxplot shows quantification of break coverage at ± 500bp of H3K27ac CHIP-seq summits (**** $p$ ~ 0, Wilcoxon signed-rand test). **G.** Cumulative, read normalized, single nucleotide resolution profile of DSBs at strong H3K4me3 sites (top 10%, n = 2845, left) and at weak H3K4me3 sites (bottom 10%, n = 2845, right) in WT and DHX36 KO cells. **H.** Boxplot shows quantification of break coverage at ± 500bp of H3K4me3 ChIP-seq peaks (**** $p$ ~ 0, Wilcoxon signed-rand test). **I.** Cumulative, read normalized, single nucleotide resolution profile of DSBs at strong CTCF-binding sites (top 10%, n = 6820, left) and at weak CTCF-binding sites (bottom 10%, n = 6820, right) in WT and DHX36 KO cell lines. **J.** Boxplot shows quantification of break coverage at ± 150bp of CTCF ChIP-seq peaks (**** $p$ ~ 0, Wilcoxon signed-rand test). RPM – reads per million.

### Figure S6

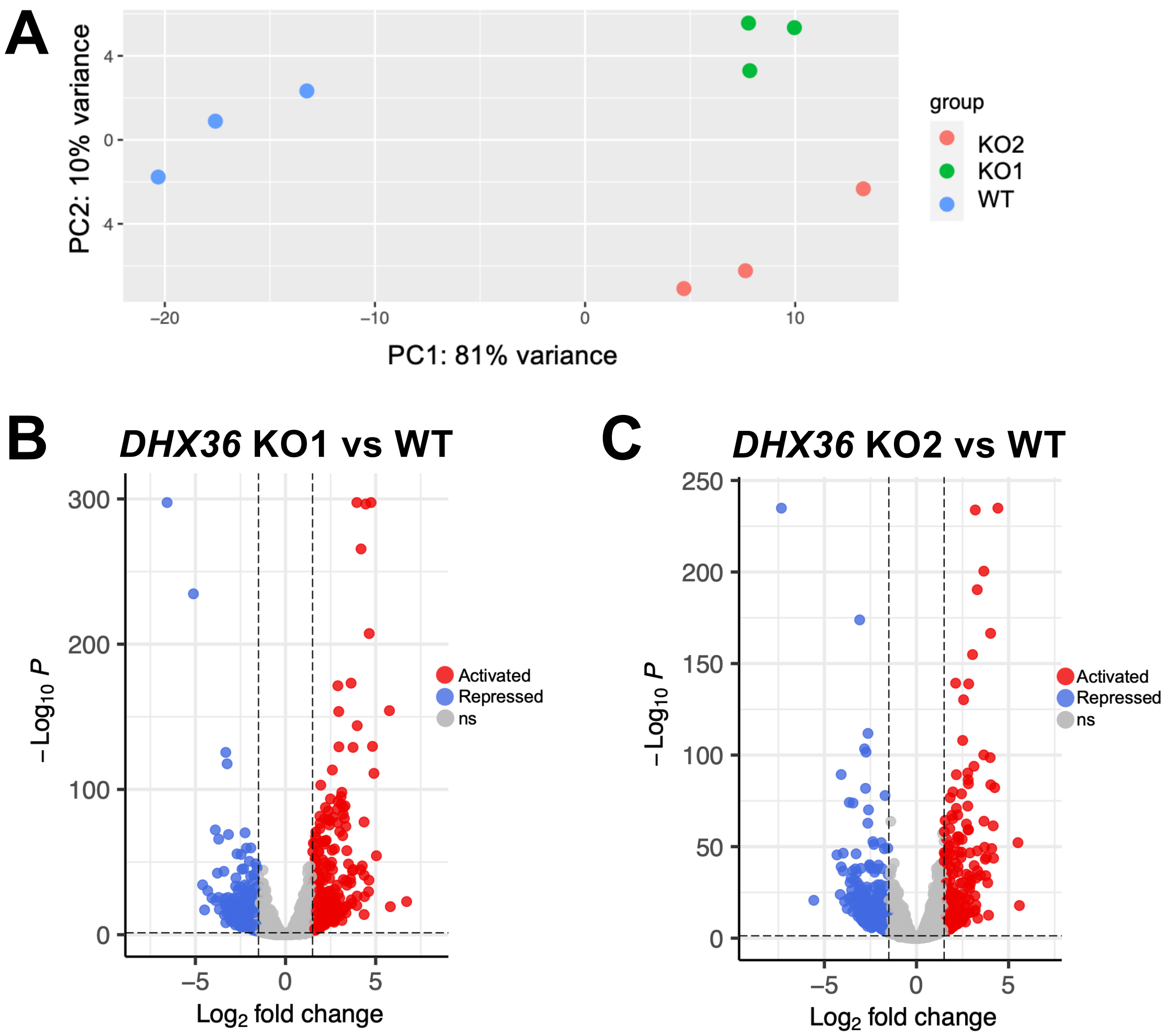


**Figure S6. Depletion of DHX36 helicases changes global gene expression in Jurkat cells. A.** Principal component analysis shows clustering of the Jurkat WT and DHX36 KO cell lines. Differentially expressed genes in DHX36 KO1 (**B**) and KO2 (**C**), when compared to the WT (red – activated genes, blue – repressed genes, grey – ns, not significantly changed genes; total n = 15081).

### Figure S7

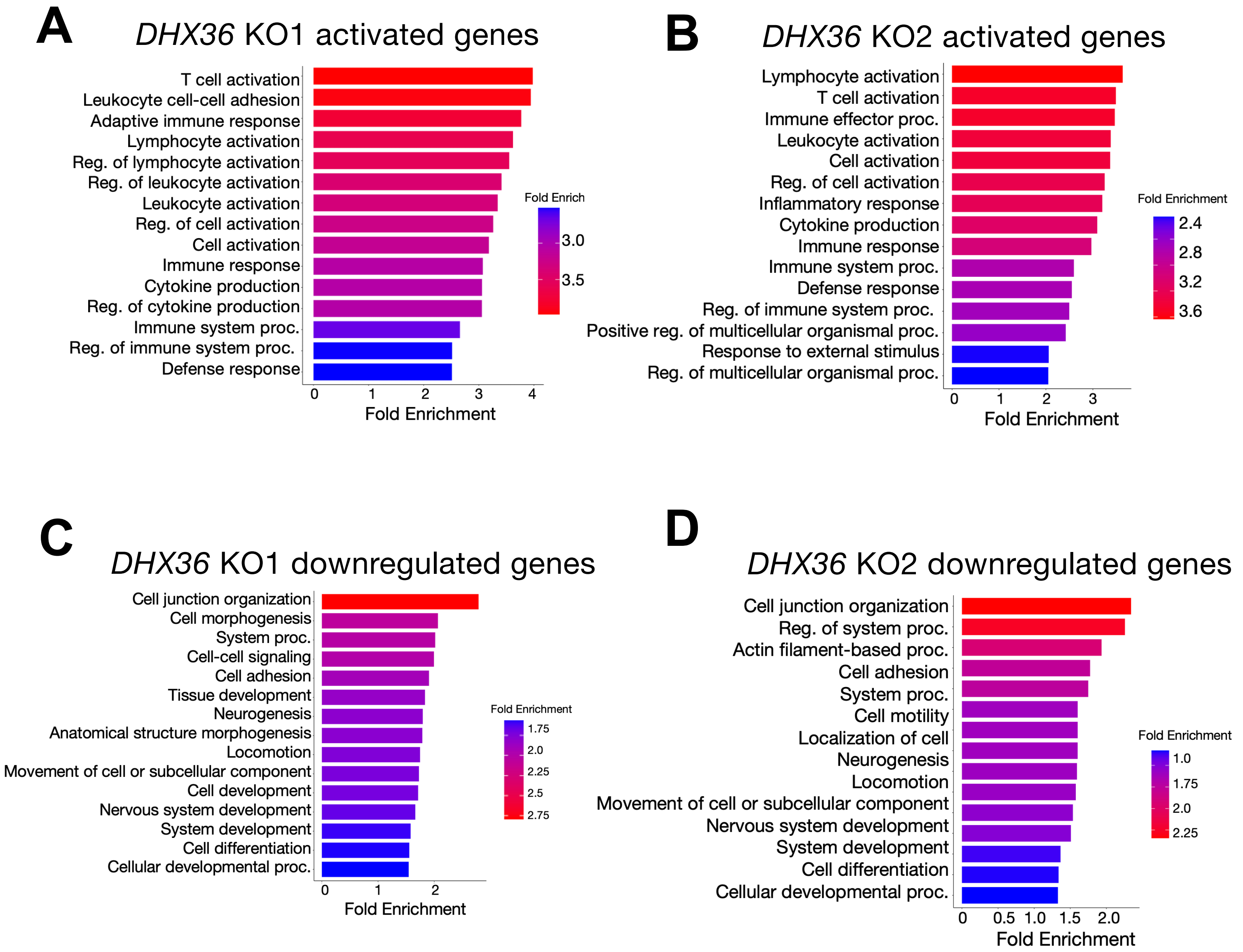


**Figure S7. Upon loss of DHX36, significantly upregulated genes are immune related, whereas the significantly downregulated genes are involved in cell junction organization and migration.** GO:Biological processes enriched in genes activated in DHX36 KO1 (n = 686) (**A**) and in DHX36 KO2 (n = 583) (**B**); Processes enriched in genes repressed in DHX36 KO1 (n = 943) (**C**) and in DHX36 KO2 (n = 1020) (**D**).

### Figure S8

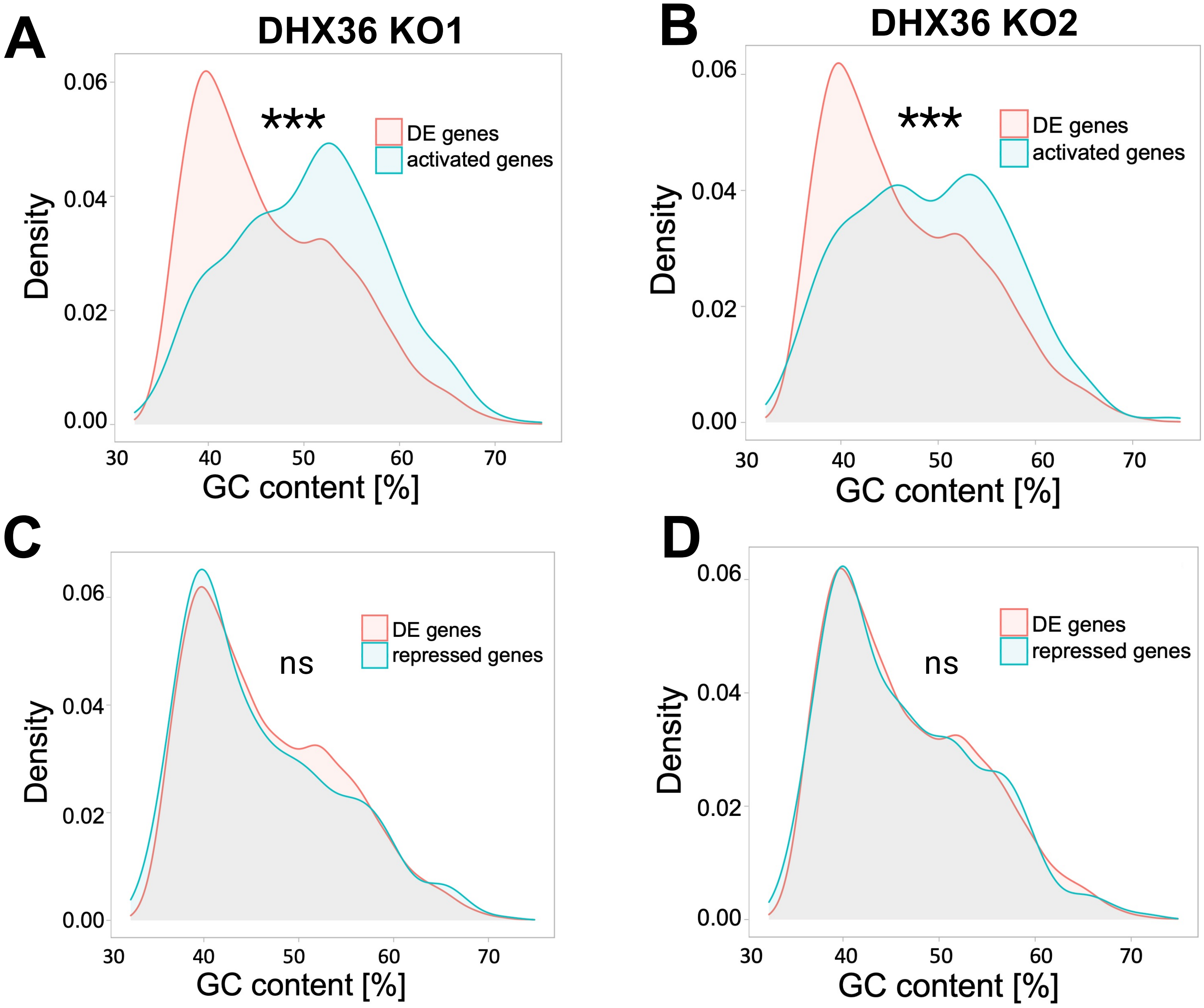


**Figure S8. Activated genes upon loss of DHX36 helicase, but not downregulated genes, show GC enrichment.** Density plot showing enrichment of GC content in upregulated genes in DHX36 KO1 (**A**) and DHX36KO2 (**B**) and in downregulated genes in DHX36 KO1 (**C**) and DHX36 KO2 (**D**) compared to all differentially expressed genes (as a background, n=15 081). *** indicates $p < 0.001$, ns – not significant, Chi-squared test.

### Figure S9

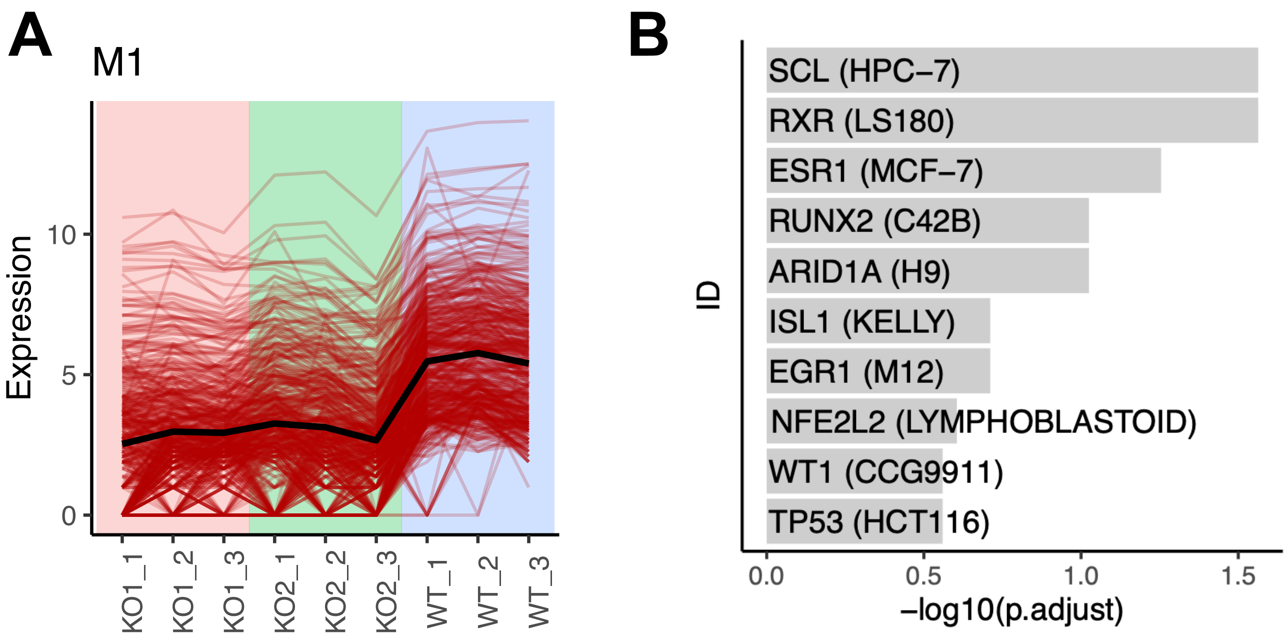


**Figure S9. Loss of DHX36 helicase leads to changes in Module M1 expression. A.** Expression profile of 465 genes in Module M1 that are downregulated in the DHX36 KO cell lines compared to the WT. **B.** Genes in M1 are enriched for targets of SCL and RXR.

### Figure S10

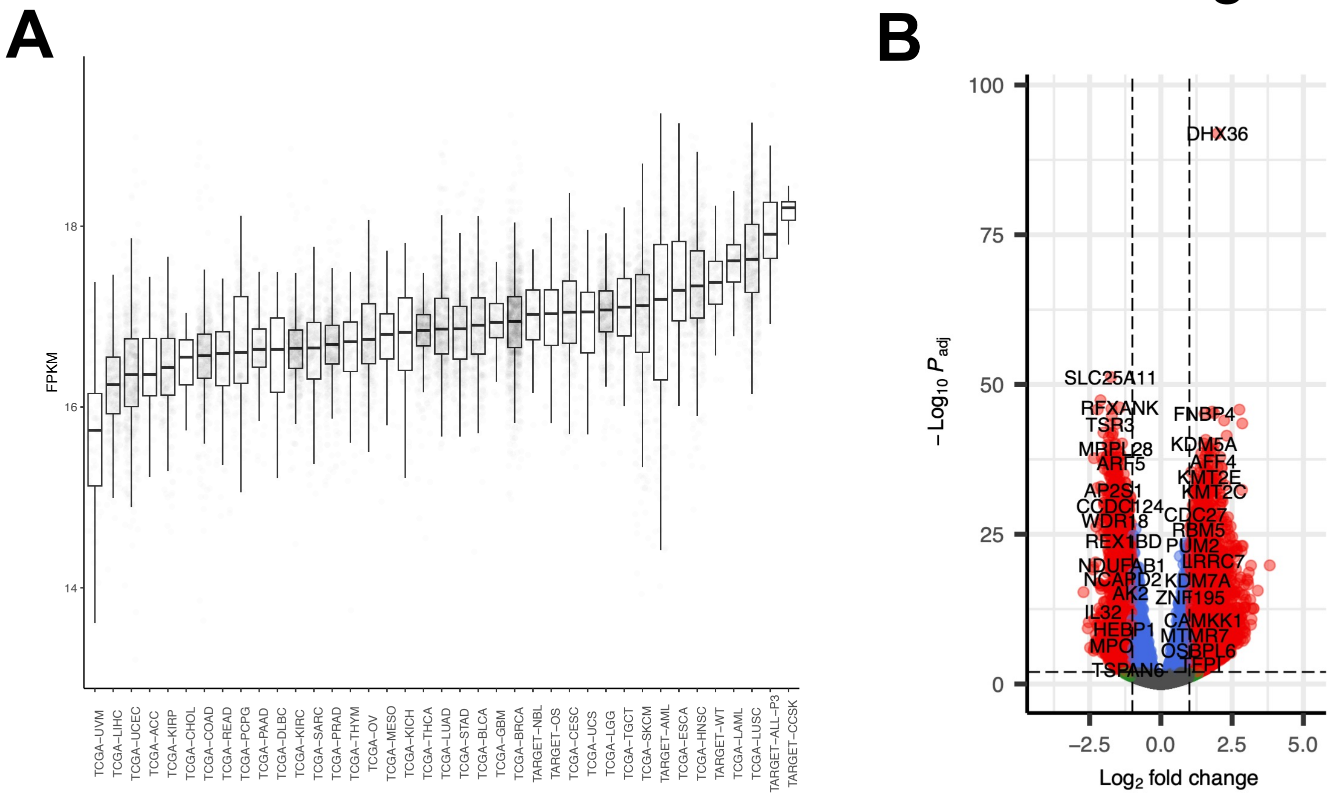


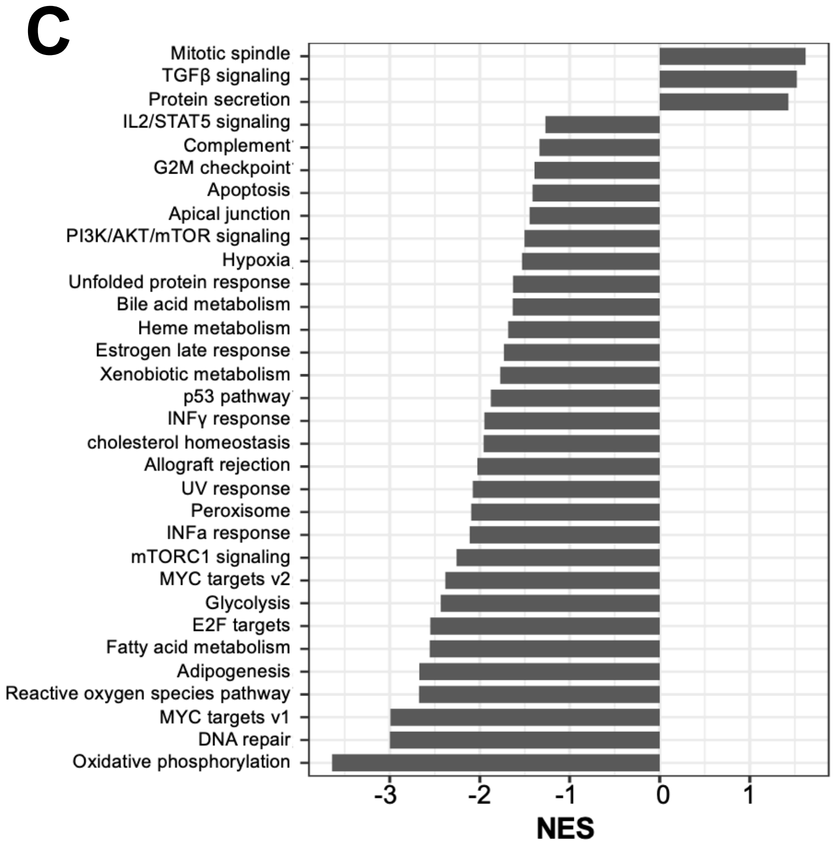


**Figure S10. DHX36 expression profile in B-ALL patients. A.** DHX36 mRNA expression profiled in 40 cancers from TCGA and TARGET shows the highest median expression in CCSK and ALL. *DHX36* expression is highest in pediatric B-ALL and CCSK. The pan-cancer normalized expression data (log2(fpkm-uq+1)) were downloaded from UCSC Xena and grouped based on the project_id for this plot. **B.** Volcano plot of differentially expressed genes between *DHX36*-high and *DHX36*-low group for the cohort of 108 pediatric B-ALL patients. The differential gene expression analysis between DHX36-HIGH and DHX36-LOW groups were performed using DESeq2, accounting for gender and cohort differences. **C.** GSEA analysis of the molecular signature database of hallmark gene sets showed low *DHX36* expression correlation with increased interferon production and DNA repair. The GSEA analysis ranked the genes based on the Wald statistic.

### Figure S11

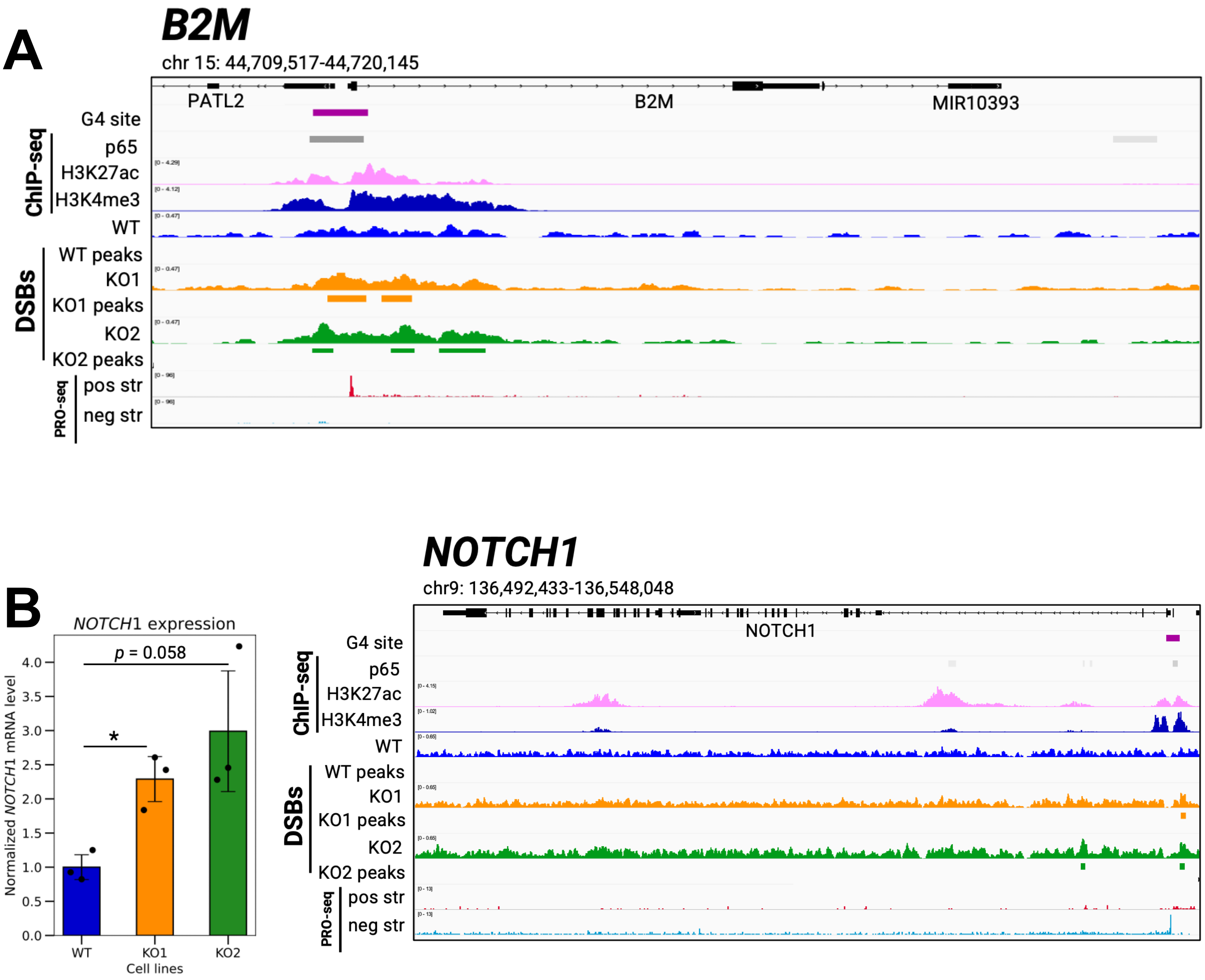


**Figure S11. Break cluster sites upon loss of DHX36.** Genome browser at *B2M* gene (**A**) and *NOTCH1* gene (**B**) showed enrichment of DBSs and break peaks upon loss of DHX36, which overlap with G4 consensus sites, p65 binding, and Pol II promoter-proximal pausing sites by PRO-seq. Expression level of *NOTCH1* is upregulated upon loss of DHX36 (* $p$ value < 0.05, Student t-test).
